# Hypomethylated poplars show higher tolerance to water deficit and highlight a dual role for DNA methylation in shoot meristem: regulation of stress response and genome integrity

**DOI:** 10.1101/2020.04.16.045328

**Authors:** M.D. Sow, A-L. Le Gac, R. Fichot, S. Lanciano, A. Delaunay, I. Le Jan, M-C. Lesage-Descauses, S. Citerne, J. Caius, V. Brunaud, L. Soubigou-Taconnat, H. Cochard, V. Segura, C. Chaparro, C. Grunau, C. Daviaud, J. Tost, F. Brignolas, S.H. Strauss, M. Mirouze, S. Maury

**Author notes:** Corresponding author, The author responsible for distribution of materials integral to the findings presented in this article in accordance with the policy described in the Instructions for Authors (www.plantcell.org) is: Stéphane Maury. Institute for Biology III, University of Freiburg, 79104 Freiburg, Germany. AGAP, Université Montpellier, CIRAD, INRA, Montpellier SupAgro, UMR 1334, 34060 Montpellier, France. First co-authors with equal contribution (M.D. Sow, A-L. Le Gac).

## Abstract

As fixed and long living organisms subjected to repeated environmental stresses, trees have developed mechanisms such as phenotypic plasticity that help them to cope with fluctuating environmental conditions. Here, we tested the role DNA methylation as a hub of integration, linking plasticity and physiological response to water deficit in the shoot apical meristem of the model tree poplar (*Populus*). Using a reverse genetic approach, we compared hypomethylated RNAi-*ddm1* lines to wild-type trees for drought tolerance. An integrative analysis was realized with phytohormone balance, methylomes, transcriptomes and mobilomes.

Hypomethylated lines were more tolerant when subjected to moderate water deficit and were intrinsically more tolerant to drought-induced cavitation. The alteration of the *DDM1* machinery induced variation in DNA methylation in a cytosine context dependent manner, both in genes and transposable elements. Hypomethylated lines subjected to water deficit showed altered expression of genes involved in phytohormone pathways, such as salicylic acid and modified hormonal balance. Several transposable elements showed stress- and/or line-specific patterns of reactivation, and we could detect copy number variations for two of them in stressed *ddm1* lines.

Overall, our data highlight two major roles for DNA methylation in the shoot apical meristem: control of stress response and plasticity through transduction of hormone signaling and maintenance of genome integrity through the control of transposable elements.

## INTRODUCTION

As long living organisms, trees are subjected to repeated environmental challenges over their lifetime. In recent decades, forest stresses have grown; forest decline has been reported around the world due to heat and drought stress episodes (Allen et al., 2010; Anderegg et al., 2016). In order to survive, they can adjust rapidly using epigenetic as well as physiological levels of regulation (Nicotra et al., 2015). Epigenetics is defined as the study of heritable changes that affect gene expression without changing the DNA sequence (Russo et al., 1996). Efforts have been made, primarily in annuals, to unravel the role of epigenetic mechanisms (in particular DNA methylation) in plant developmental processes, stress response, plasticity, and adaptation (Slotkin and Martienssen, 2007; Colomé-Tatché et al., 2012; Cortijo et al., 2014; Kooke et al., 2015; Raju et al., 2018; Schmid et al., 2018).

Although epigenetic processes are commonly regarded as a source of flexibility in perennials like trees (Bräutigam et al., 2013; Zhu et al., 2013; Yakovlev et al., 2012, 2016; Carneros et al., 2017; Plomion et al., 2016; Lafon-Placette et al., 2018; Sow et al., 2018b), the functional role of DNA methylation in forest trees under environmental changes is still unclear. As a model tree with important genomic resources (Tuskan et al., 2006; Jansson and Douglas, 2007), poplar (*Populus* spp.) has been a prime system for the study of the ecophysiological and molecular basis of drought response (Monclus et al., 2006; Street et al., 2006; Bogeat Triboulot et al., 2007; Cohen et al., 2010; Hamanishi et al., 2012; Fichot et al., 2015). For example, differences in global DNA methylation levels among poplar hybrid genotypes have been shown to correlate with biomass production under water deficit (Gourcilleau et al., 2010; Raj et al., 2011; Le Gac et al., 2019). Recently, DNA methylation-based models have been proposed as a strategy to validate the identity, provenance or quality of agro-forestry products (Champigny et al., 2019). Lafon-Placette et al. (2018) established that drought preferentially induced changes in the DNA methylation of phytohormone-related genes, apparently elevating phenotypic plasticity (Lafon-Placette et al., 2018). This has raised the question of a possible link between epigenetics and phytohormone signaling / synthesis in order to explain plasticity in plants, in particular in meristematic tissues where development takes place (Maury et al., 2019). Phytohormones are key regulatory elements in plant development and stress response, and their action is often fast and transient. Epigenetic regulations could play a role by modifying the expression of genes in the phytohormone pathways, notably by maintaining them after a hormonal peak, or priming their expression through time to remember the stress episode (i.e., epigenetic memory). In line with this, it has been shown that winter-dormant shoot apical meristems of poplar genotypes grown in field conditions can keep an epigenetic memory of a summer drought episode experienced during the growing season through modifications in DNA methylation (Le Gac et al., 2018; Sow et al., 2018a). The role of epigenetic memory in trees besides poplar, in response to biotic and abiotic stresses, is becoming increasingly documented (Yakovlev et al., 2014; Carneros et al., 2017; Gömöry et al., 2017; Yakovlev and Fosdal, 2017).

In plants, DNA methylation occurs in three different contexts (CHH, CHG and CpG, with H=A, C or T), and the methylation or demethylation of cytosines (*de novo* or maintenance during replication) is ensured by different DNA methyltransferases or DNA glycosylases/lyases, respectively (Zemach et al., 2010; Zhang et al., 2018). DNA methylation affects gene expression (Niederhuth and Schmitz, 2017; Bewick and Schmitz, 2017). While methylation in promoters is usually associated with gene silencing, gene-body methylation is more complex, and is often linked to tissue-specific expression or alternative splicing (Vining et al., 2012; Lafon-Placette et al., 2013; Maor et al., 2015; Zhu et al., 2016; Zhu et al., 2018).

So far, most of the studies conducted on trees and focusing on DNA methylation and gene expression have used a correlative approach. In poplar, extensive gene-body methylation is found in the open chromatin state, and is linked to structural gene characteristics, and correlated with tissue-specific gene expression or stress (Vining et al., 2012; Bräutigam et al., 2013; Lafon-Placette et al., 2013; Liang et al., 2014; Lafon-Placette et al., 2018). In addition to controlling gene expression, it is clear that methylation also helps to maintain genome integrity by silencing the relics of viral genomes (i.e., transposable elements, TEs), stopping them from spreading within the host genome (Ikeda and Nishimura, 2015; Fultz et al., 2015). For a long time considered as ‘junk DNA,’ the evolutionary impact of TEs is now well established, and TEs contribute strongly to genome plasticity in eukaryotes (Lisch, 2012; Ayarpadikannan and Kim, 2014; Hirsch and Springer, 2017). In plants, most TEs can be activated, and their contribution to genome size is not negligible, especially for trees with large genomes (Kejnovsky et al., 2012; Lee and Kim, 2014). DNA methylation is required to silence these TEs located in the heterochromatin, and a decrease in DNA methylation level could result in their reactivation (Lippman et al., 2004; Mirouze et al., 2009; Mirouze and Paszkowski, 2011). The overall functional role of DNA methylation, both for control of gene expression and TE dynamics in a developmental and ecological context, is poorly understood.

*DDM1* belongs to the SWI/SNF chromatin remodeling complex, and encodes a chromatin remodeling factor required for the maintenance of DNA methylation. Its depletion affects the distribution of methylation in all sequence contexts (Vongs et al., 1993; Jeddeloh et al., 1998; Gendrel et al., 2002; Zhu et al., 2013; Zemach et al., 2013). *DDM1* was first identified in *Arabidopsis* through EMS (ethyl methane sulfonate) treatment, which caused a “decrease in DNA methylation” (Vongs et al., 1993). Vongs et al., showed that *Arabidopsis ddm1* mutants displayed a 70 to 75% reduction in cytosine methylation compared to the wild-type (WT). Nonetheless phenotypic variations only appeared several generations after the initial loss of *DDM1* activity, notably through reactivation of TEs (Miura et al., 2001). Several studies further characterized *ddm1* mutants in Arabidopsis (Saze and Kakutani, 2007; Yao et al., 2012; Zemach et al., 2013; Cortijo et al., 2014; Ito et al., 2015; Kawanabe et al., 2016), turnip (Fujimoto et al., 2008; Sassaki et al., 2011), maize (Li et al., 2014), and rice (Higo et al., 2012; Tan et al., 2016). In poplar, RNAi *ddm1* lines have been obtained by targeting the transcripts of the two orthologous *DDM1* paralogs in *Populus tremula × Populus alba* cv. INRA 717-1B4 (Zhu et al., 2013). Under standard greenhouse conditions, the regenerated lines did not show developmental defects, but newly formed leaves displayed a mottled phenotype after a cycle of dormancy (Zhu et al., 2013). These RNAi *ddm1* lines have never been studied under stressed conditions.

To investigate whether variation in DNA methylation has the potential to facilitate tree plasticity under stress, we studied these *ddm1* RNAi lines during a water stress experiment. Here, plasticity is defined as modifications in phenotype (growth, morphology, anatomy, or gene expression) under a drought – re-watering regime for a given genotype (Bradshaw, 2006; Nicotra et al., 2010, 2015). To address this question in a developmental context, we focused our analysis on the shoot apical meristem (SAM, center of plant morphogenesis) one week after re-watering in order to focus on ‘stable’ post-stress epigenetic events (Lafon-Placette et al., 2018). Previous studies have shown that SAM is a critical organ where epigenetic modifications can affect plant development (Gourcilleau et al., 2010; Lafon-Placette et al., 2013; Conde et al., 2017; Lafon-Placette et al., 2018; Le Gac et al., 2018, Sow et al., 2018; Maury et al., 2019). To examine whether plasticity was associated with epigenetic variation within wild type or RNAi lines, we combined a fine scale ecophysiological characterization of growth dynamics and water relations with genomics (identification of differentially methylated regions (DMR) using whole genome bisulfite sequencing, differentially expressed genes (DEG) using RNA-seq, and active transposable elements using mobilome-seq). We report a comprehensive analysis of the functional role of DNA methylation in the poplar SAM in terms of gene expression, reactivation of TEs, and hormonal balance in response to variations in water availability.

## RESULTS

### Phenotypic and physiological differences among lines under well-watered conditions

Plants from the WW treatment remained close to field capacity during the whole experiment, with relative extractable water (REW) never dropping below 70% (Fig. 1) and Ψ_pd_ values remaining above -0.5 MPa (Sup. Fig. 2A). There was no significant difference in either REW or Ψ_pd_ among lines (Fig. 1 and Sup. Fig. 2A). The WT and *ddm1* lines all showed linear growth, and exhibited a similar height growth rate during the experiment (1.27 ± 0.07 cm.day^-1^, *P* = 0.67; Fig. 2A). However, the WT had a slightly higher diameter growth rate than the *ddm1* lines (0.11 ± 0.02 *vs.* 0.08 ± 0.01 mm.day^-1^, *P* = 0.036; Fig. 2A). Differences among lines were much stronger for total leaf area, with *ddm1* lines exhibiting significantly lower values (28 % reduction on average, *P* = 0.021) compared to the WT (Sup. Fig. 3). This was explained by a particular vertical profile of individual leaf area in the middle canopy (Sup. Fig. 3).

**Figure 1:**
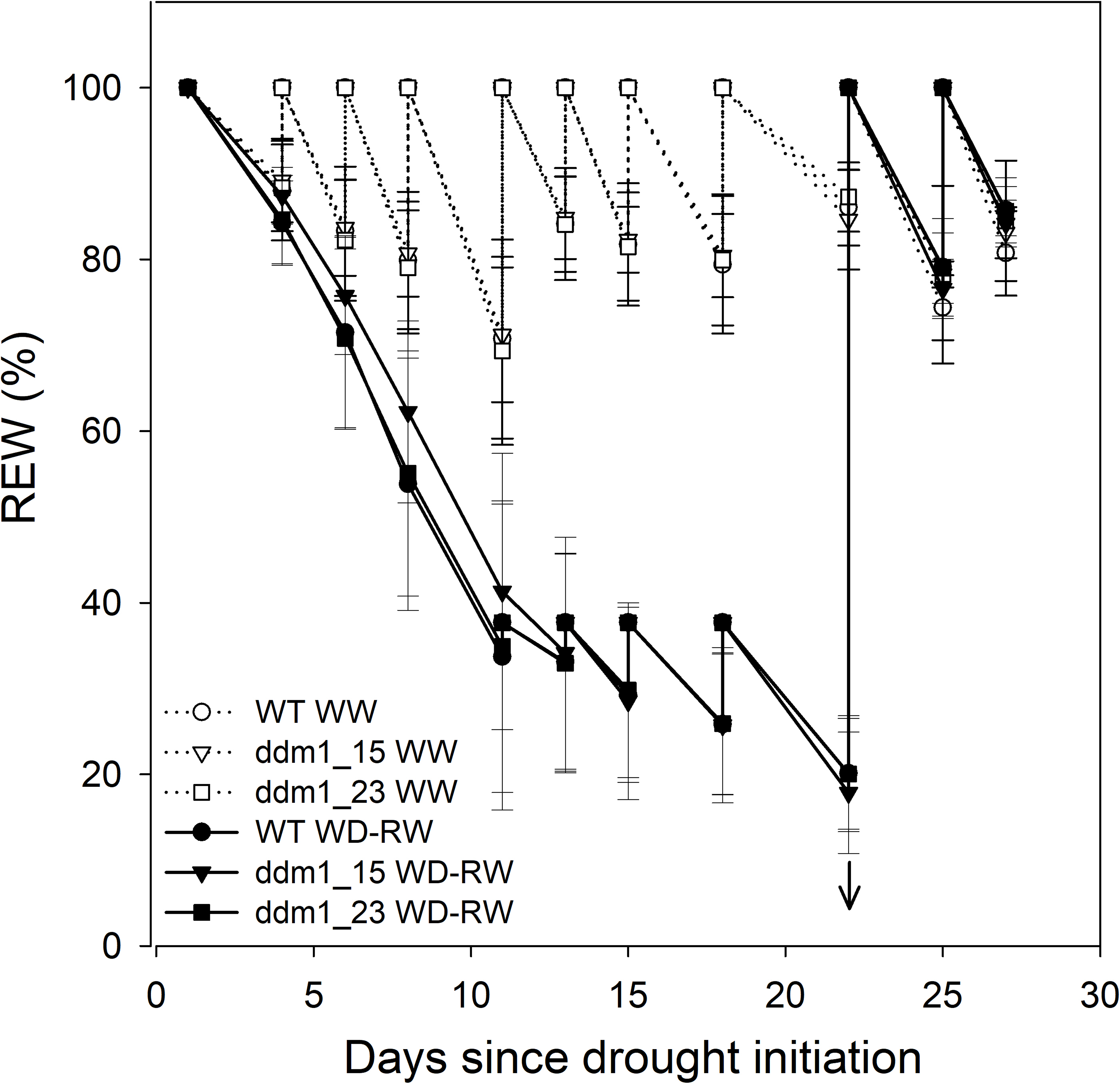
Time course of soil relative extractable water (REW) during the experiment for the wild type and the two RNAi-*ddm1* (*ddm1*-15, *ddm1*-23) poplar lines in control (well-watered, WW) and stress (moderate water deficit followed by rewatering, WD-RW) treatments. Open symbols and dashed lines for WW treatment; closed symbols and solid lines for WD-RW treatment. Circles for the wild type; triangles for the RNAi-*ddm1*-15 line; squares for the RNAi-*ddm1*-23 line. The arrows represent the end of the water deficit and the onset of rewatering. Values are genotypic means ± SE (*n* = 6 per line for WW, *n* = 12 per line for WD-RW).

**Figure 2:**
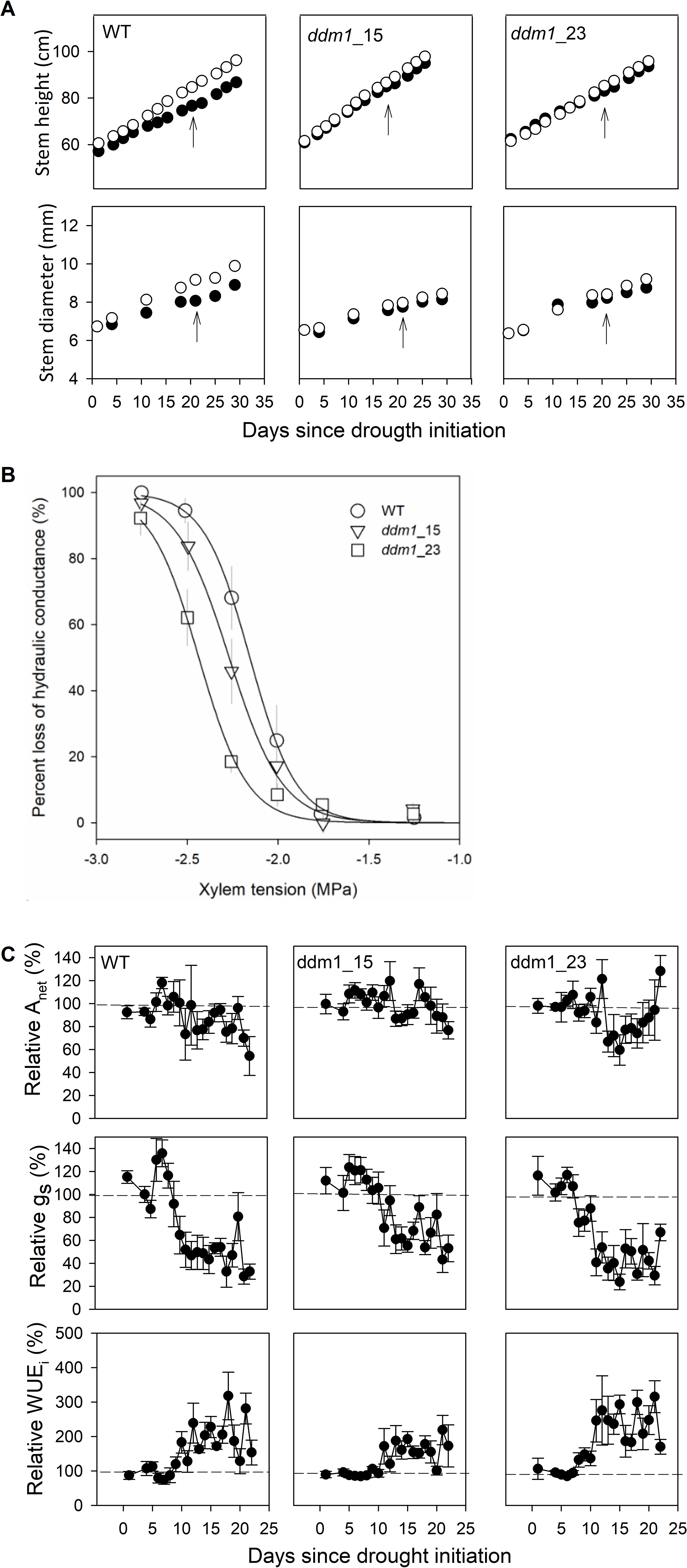
Phenotypic and physiological characterization of the wild type and the two RNAi-*ddm1* (*ddm1*-15, *ddm1*-23) poplar lines in control (well-watered, WW) and stress (moderate water deficit followed by rewatering, WD-RW) treatments. Open symbols and open bars for WW; closed symbols and closed bars for WD-RW. Values are genotypic means ± SE. **A.** Time course of stem height and diameter (*n* = 6 per line in WW, *n* = 12 per line in WD-RW). The arrows represent the end of the water deficit and the onset of rewatering. **B**. Xylem vulnerability to drought-induced cavitation measured at the end of the experiment (*n* = 6 per line). **C**. Time course of leaf gas exchange (A_net_, net CO_2_ assimilation rate, g_s_, stomatal conductance to water vapour, WUE_i_, intrinsic water-use efficiency computed as A_net_/g_s_). Values presented are those of WD-RW plants relative to WW controls (*n* = 5 per line per treatment). Treatment effects were evaluated within each line using a t-test. Levels of significance are *, 0.01 < *P* < 0.05; **, 0.001 < *P* < 0.01; ***, *P* < 0.001; ns, non-significant.

All lines showed comparable leaf Ψ_min_, leaf δ^13^C and stomatal density (Sup. Fig. 2B), but significant differences were observed for xylem vulnerability to cavitation (*P* < 0.001, Fig. 2B). The WT was the most vulnerable (P_50_ = −2.16 ± 0.05 MPa), while *ddm1-23* was the most resistant (P_50_ = −2.45 ± 0.04 MPa), and *ddm1-15* was intermediate (P_50_ = −2.28 ± 0.04 MPa) (Table 1). Differences among lines were not found in other xylem structural or biochemical traits (Table 1).

**Table 1:**
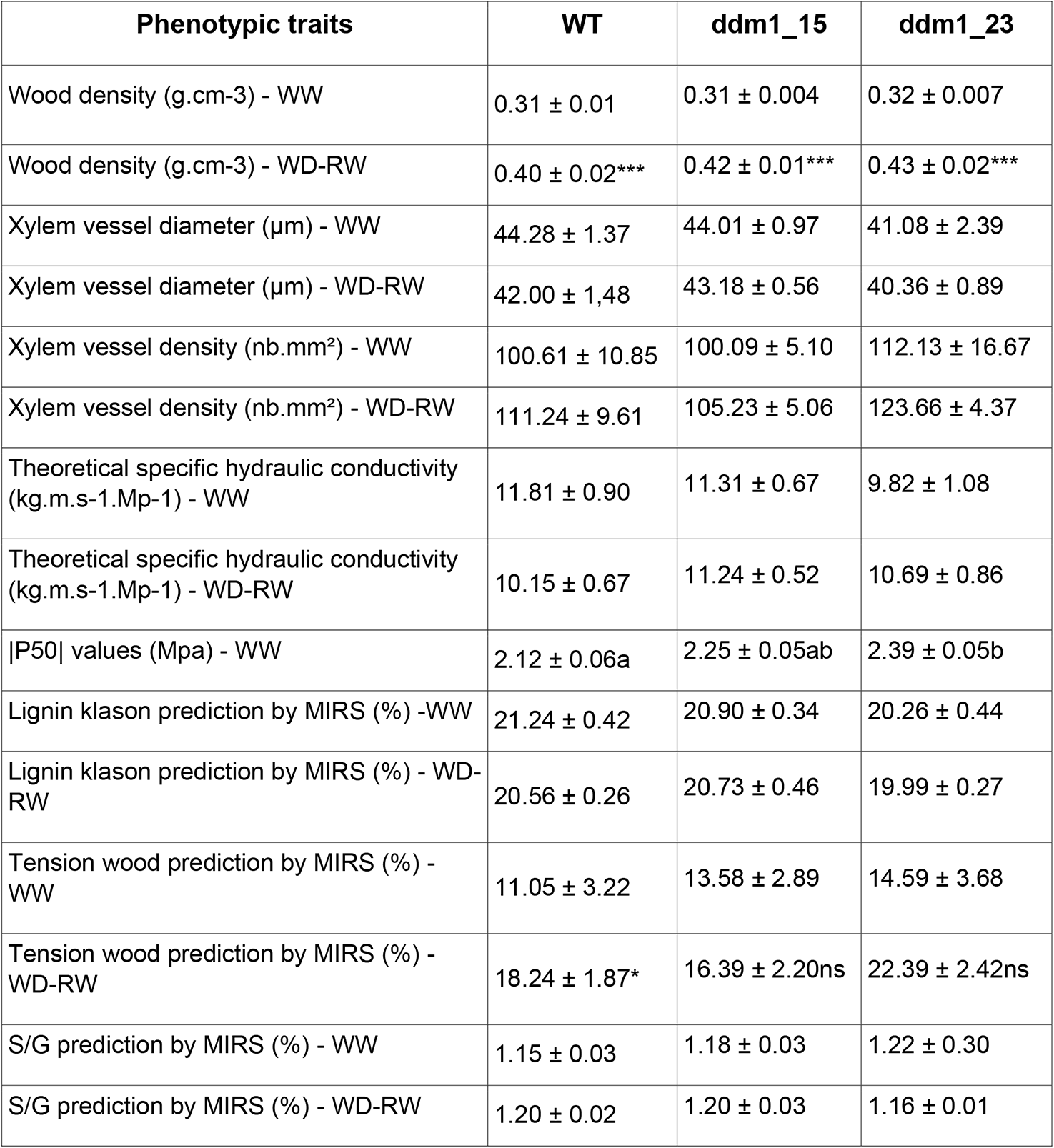
Xylem structural, functional and biochemical traits measured for the wild type and the two RNAi-*ddm1* (*ddm1*-15, *ddm1*-23) poplar lines in control (well-watered, WW) and stress (moderate water deficit followed by rewatering, WD-RW) treatments. Values are genotypic means ± SE (*n* = 6 per line per treatment). The P_50_ is the xylem tension inducing 50% loss of hydraulic conductance estimated from vulnerability curves (see Materials and Methods for additional information). S/G corresponds to the ratio between syringyl-like (S) and guaiacyl-like lignin monomeric units (G). Treatment effects were evaluated within each line by using a t-test. Different letters indicate significant differences between genotypes within treatments following Tukey’s post hoc test. Levels of significance are *, 0.01 < *P* < 0.05; **, 0.001 < *P* < 0.01; ***, *P* < 0.001; ns, non-significant; na, not available.

The proportion of “mottled leaves” reached 40% for *ddm1-23,* and more than 60% for *ddm1-15* at the end of the experiment, while it remained close to zero for the WT (Sup. Fig. 4). Symptom occurrence was not linear but tended to increase at a specific physiological stage (Sup. Fig. 4). In addition, the line *ddm1-23* exhibited several leaves that tended to fold around the midvein, a physical defect that was not found in the WT, and only rarely in *ddm1-15* (Sup. Fig. 4).

### Differences in drought response

Soil water content of plants in drought conditions started to be significantly lower four days after the initiation of water deficit. Values of REW were maintained around 35% until t_1_; re-watering at t_1_ increased REW back to control values (Fig. 1). Predawn leaf water potential (Ψ_pd_) at t_1_ was significantly lower than in well-watered plants (*P* < 0.001), and reached approximately -0.8 MPa, with no difference among lines (Sup. Fig. 2A); in contrast, Ψ_min_ values were not significantly affected by drought (Sup. Fig. 2A). Height and diameter growth rates during drought were significantly lower in the WT only (Fig. 2A, *P* < 0.001 and *P* = 0.026, respectively). The effect persisted after re-watering until t_2_ for diameter (*P* < 0.001) while height growth recovered to control values.

In response to water deficit, g_s_ started to decrease significantly 10 days after drought initiation, *i.e.* once REW had dropped below 40% (*P* = 0.027 for WT, *P* = 0,171 for ddm1_15 and *P* = 0.021 for ddm1_23 Fig. 1, 2C). The WT and *ddm1-23* showed relatively comparable dynamics, and reached almost an 80% decrease relative to controls, while stomatal closure was less pronounced in *ddm1-15* (Fig. 2C). A_net_ was less impacted than g_s_, especially in *ddm1-15*, in agreement with the moderate intensity of water deficit (Fig. 2C). Therefore, the increase in intrinsic water-use efficiency (*i.e.* the ratio A_net_/g_s_) was more pronounced in the WT and *ddm1-23* (Fig. 2C), which was confirmed by a greater δ^13^C increase in these same lines (Sup. Fig. 2B). Total stomatal density was not impacted by water deficit in *ddm1* lines, while it was significantly increased in the WT (*P* = 0.030, Sup. Fig. 2C). Xylem traits were only seldom affected by water deficit (Table 1). Xylem density was significantly increased in all lines (*P* < 0.001), while the proportion of tension wood increased in the WT only (Table 1). Water deficit had no significant effect on the occurrence of mottled or folded leaves (Sup. Fig. 4).

### Phytohormone concentration in SAMs

There was no significant change among lines in terms of salicylic acid (SA) or the different types of cytokinin in the WW treatment (Fig. 3). In response to water deficit-rewatering, SA increased while zeatin riboside and zeatin -O-glucoside riboside decreased in RNAi lines compared to the WT (Fig. 3). The other type of cytokinin, the isopentenyladenosine, showed a more complex pattern in RNAi lines. In addition, the ddm1-23 line showed a significant response for SA and for two cytokinins, with a similar trend to WT (Fig. 3). The *ddm1-15* line showed a significant response for one cytokinin only (Fig. 3). ABA and free auxin remained unaffected in all treatments.

**Figure 3:**
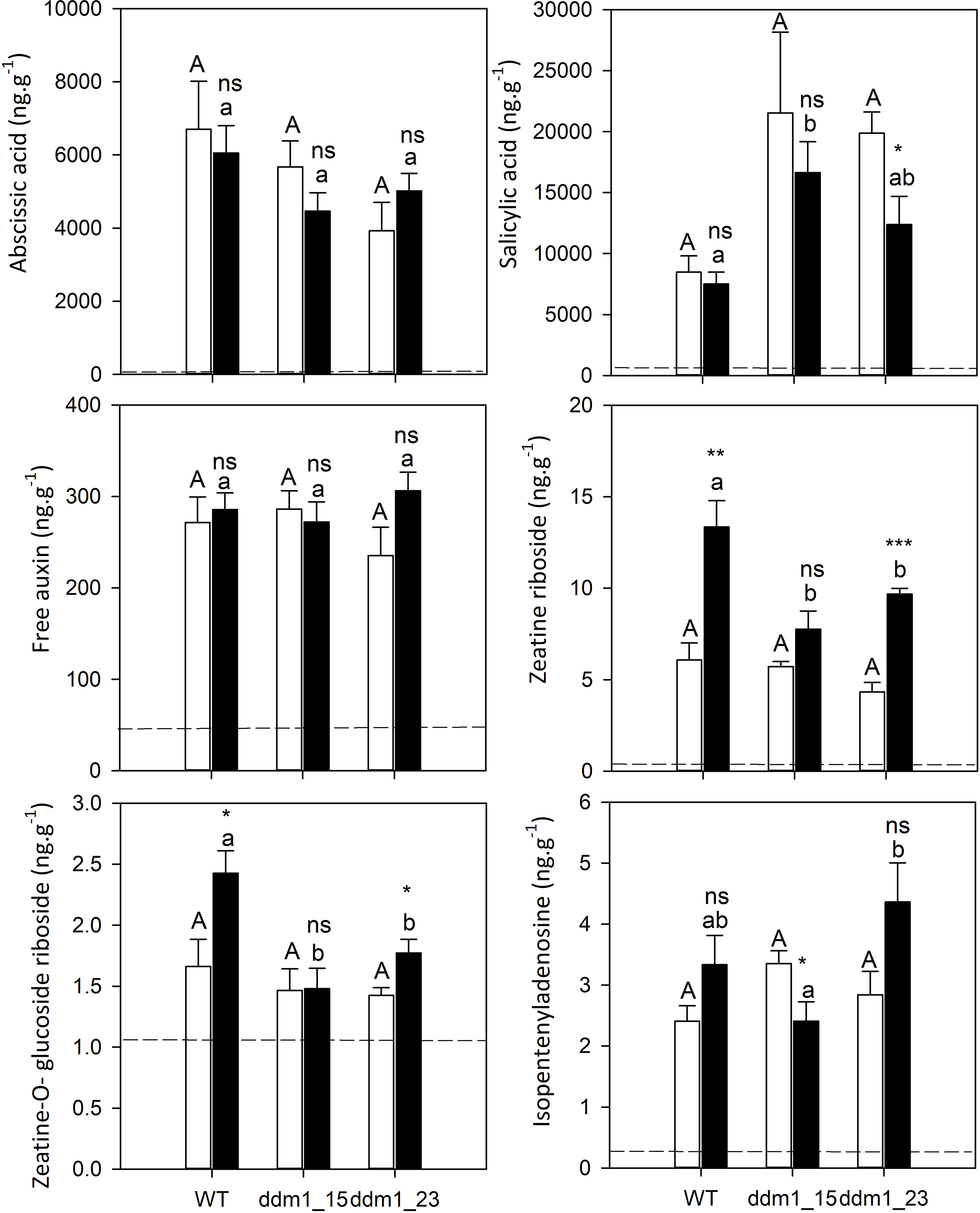
Phytohormone contents (cytokinins, abscisic acid, salicylic acid and free auxin) measured in shoot apex just after rewatering (t_1_). Values are genotypic means ± SE (*n* = 3 per line in WW and in WD-RW). Treatment effects were evaluated within each line by using t-test. Levels of significance are *: p< 0.05, **: p< 0.01, ***: p< 0.001 and ns: non-significant. The different letters indicate the differences between lines within a water regime following a Tukey’s post hoc test (lower case letters for WW, capital letters for WD-RW).

### Methylome analysis and identification of line- and stress-specific DMRs

Global DNA methylation content (HPLC analysis) ranged from 17.5 to 21.3 % between lines and treatments (Sup. Fig. 5A). There was no significant line × treatment interaction effect. Global DNA methylation was significantly lower in RNAi lines than in the WT under water deficit only, although there was no significant effect of water deficit (Sup. Fig. 5A). Cytosine methylation percentages (WGBS analysis) for the three contexts ranged from 18.6 to 19.6% in CpG, 4.4 to 6.0 in CHG and 1.6 to 2.0 in CHH contexts, with the *ddm1-15* lines displaying the lowest values in all contexts (Sup. Table 1, Sup. Fig. 1D).

Only a few stress-specific DMRs were identified. In contrast, thousands of line-specific DMRs were identified, especially for *ddm1-15* (Sup. Fig. 5B). The two RNAi lines displayed a different number of DMRs (29785 *vs*. 30925 for *ddm1-15* in WW and WD-RW treatments, respectively, and 11409 *vs*. 11104 for *ddm1-23* in WW and WD-RW treatments, respectively). Most of these DMRs were hypomethylated and context-dependent with higher values found in CHG context, especially in *ddm1-15* (20310 *vs*. 20847 DMRs in WW and WD-RW treatments, respectively) (Sup. Fig. 5C & 5D). Line-specific DMRs were unequally distributed in the three different contexts. In the CpG context, DMRs presented a bimodal distribution with both hypo and hypermethylated DMRs (Sup. Fig. 5E). In the CHG context, most DMRs were hypomethylated, while in the CHH context DMRs were mostly slightly hypomethylated (Sup. Fig. 5E). The number of stress-specific DMRs was similar between the WT and *ddm1* lines, and between the hypo and hypermethylated states (Fig. 4A). These DMRs were mainly found in CG and CHG contexts (Fig. 4A).

**Figure 4:**
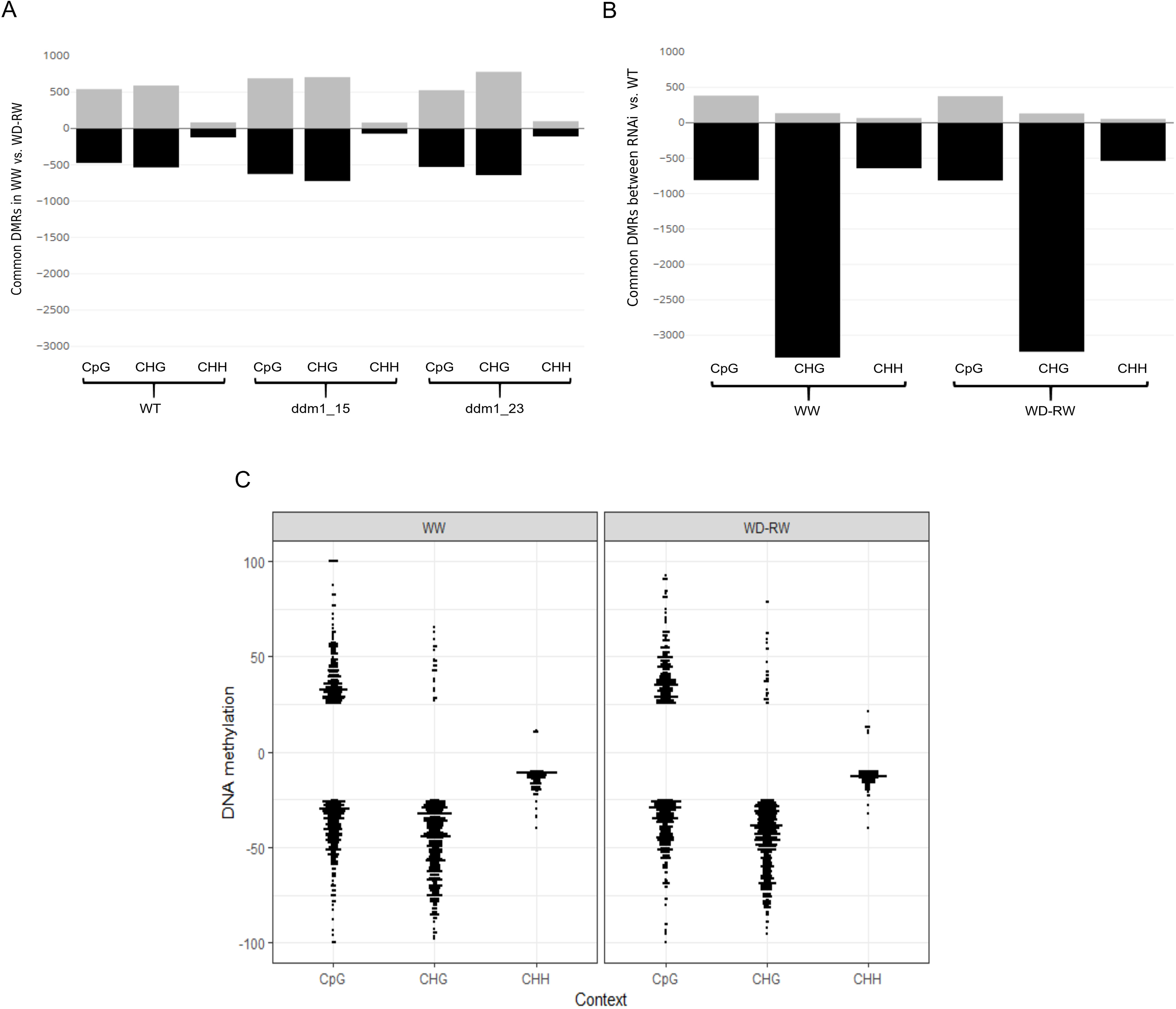
Variations in DNA methylation among RNAi and WT lines in shoot apical meristem one week after rewatering (t2). **A**. Common DMRs between WW and WD-RW conditions. Black bars for hypomethylated DMRs and grey bars for hypermethylated DMRs. **B**. Common DMRs between the two RNAi lines (ddm1_15 & ddm1_23) vs. the WT lines in all contexts (CG, CHG & CHH) and in WW and WD-RW conditions. Black bars for hypomethylated DMRs and grey bars for hypermethylated DMRs. **C**. DNA methylation variation of the common DMRs in the RNAi lines vs. the WT line in WW and WD-RW per context of methylation. Only DMRs with at least a 25 % difference were kept except for CHH where a threshold of 10% was applied due to the low proportion of DMRs in that context.

In order to focus on DMRs systematically associated with *DDM1* defects, we identified DMRs commonly shared between *ddm1-23* and *ddm1-15* RNAi lines in either WW or WD-RW treatments (Fig. 4B). Regardless of the treatment, common DMRs were systematically over-represented in the CHG context (3463 *vs*. 3378 DMRs in WW and WD-RW treatments, respectively) and were mostly hypomethylated (Fig. 4B). Among common DMRs, 19 and 21% were localized in genes, 7 and 8% in promoters (+/-2kb from the TSS) and 1% in transposable elements (TEs), for WW and WD-RW treatments, respectively (Sup. Fig. 6A). The majority of DMRs (73 and 71% in WW and WD-RW treatments, respectively) were therefore localized in intergenic regions (Sup. Fig. 6A). In the WW treatment, 879 genes were found to be strictly included within the common DMRs (hereafter called DMGs for Differentially Methylated Genes), while 910 DMGs were found in the WD-RW treatment. These numbers increased considerably (up to more than 13,000 genes) when enlarging the windows for DMR identification from 2 kb to 25 kb (Sup. Fig. 6B). In both treatments, a similar number of hypo and hypermethylated DMGs was found in the CG context, while in the CHG and CHH contexts, DMGs were mostly hypomethylated (Fig. 4C). In the CHH context, DMGs were slightly methylated (between 25 to 50% of difference), compared to the CG and CHG contexts (Fig. 4C). Gene Ontology annotation of DMGs revealed significant enrichment in biological functions such as multicellular organism development (including shoot system development), negative regulation of biological processes, and response to abiotic stress (including response to hormones) (Sup. Fig. 6C).

### Transcriptome analysis for the water deficit – rewatering condition

The identification of DEGs focused on the WD-RW treatment, but revealed clear differences between the WT and the *ddm1*-23 lines. An average of 96 % mapping efficiency was found for the *P. tremula × alba* reference genome (v1). A total of 32 048 genes were analyzed, but only 136 genes were significantly differentially expressed (*P* < 0.05). Gene ontology annotation revealed significant enrichment in functions such as defense response, including immune response, response to hormones (SA, Ethylene), response to chitin, and regulation of RNA metabolism (Fig. 5A). The 136 DEGs (76 up-regulated and 60 down-regulated) were grouped according to *Arabidopsis thaliana* gene annotation homology into seven main classes: cell death, defense response and cell wall, immune response, metabolism, signalization, transcription factors, and unknown function (Fig. 5B). Genes related to immune response were systematically up-regulated (*RBOHD, CYP94B1, RLP1, RLP56, RPM1, PLDGAMMA1, PDF1*). Most genes related to transcription factors (15/17) were also up-regulated (*WRKY, MYB106, ERF, SZF2, PDF2, SVP/AGL22*), with only two genes down-regulated (*MYB48* and *DTA2*). Defense and cell wall related genes were both up-(18, including *CHITIV*, *KTI1*, *PR4*, etc. involved in plant pathogen-interaction) and down-regulated (13). Phytohormone pathways were also over-represented in distinct classes with 13 DEGs (8 up- and 5 down-regulated) directly involved in defense responsive hormone biosynthetic pathways, such as SA pathways (*SAMTs*), the jasmonic acid (JA) pathway (*OPR2*, *CYP94B1*), the ethylene (ET) pathway (*ERF1*, *ERF12*), or auxin responsive genes (*SAUR29, GH3.1, IBR3, BG1, ABCG36*), gibberellic acid (GA) synthesis (*GA3OX1*) and cytokines (CK) pathways (*AHP1*) (Fig. 5B).

**Figure 5:**
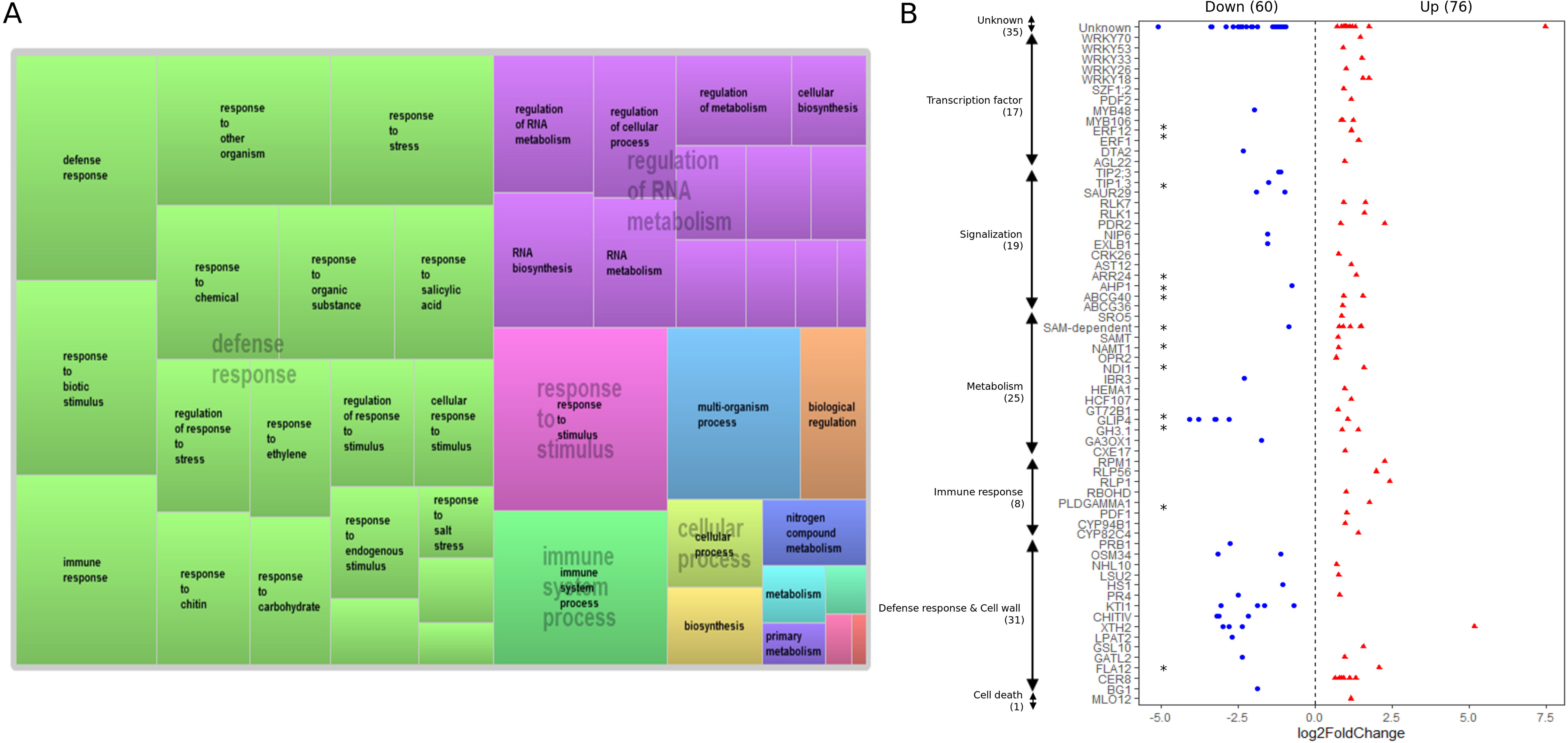
Gene expression variations in ddm1-23 vs. the WT line in non-irrigated condition (WD-RW) in shoot apical meristem collected one week after rewatering (t2). **A**. GO annotation of the differentially expressed genes (genes with an adjusted p-value by FDR, false discovery rate < 0.05 = 136 DEGs identified) between ddm1-23 and the WT line. GO labels were retrieved from popgenie and treemap realized with REVIGO. **B**. Annotation of DEGs with expression variation values (log2FoldChange) using GO labels retrieved from popgenie. Blue for downregulated genes and red for upregulated genes. The * indicates hormone related genes found in DEGs. The numbers (1), (31), (8), (25), (19), (17), (35) represent respectively the number of DEGs found in Cell death, Defense & Cell wall, Immune response, Metabolism, Signalization, Transcription factors and Unknown processes respectively. Log2FoldChange = log-ratio of normalized mean read counts in RNAi vs. WT lines.

In order to test the link between gene expression and DNA methylation, DEGs were co-localized with common DMRs among *ddm1* lines. Although only seven DEGs (Potri.001G048700, Potri.001G065300, Potri.002G192400, Potri.009G051300, Potri.016G130900, Potri.T041700, Potri.T085000) overlapped with the DMR genomic locations, 53 were located in the direct vicinity of a DMR (+/- 10 kb) and 98 at +/- 25 kb (Sup. Fig. 7A). A significant and negative rank correlation (Spearman’s *rho* = −0.32, *P* < 0.001) was observed between methylation in the three contexts and expression values when considering at least a +/- 10 kb window for DMRs to reach statistical significance (Sup. Fig. 7B).

### Mobilome analysis and identification of line- and/or stress-specific active TEs

We identified between 44 (ddm1-23 in WW) and 169 (ddm1-15 in WD-RW) TE families producing extrachromosomal circular DNAs (eccDNAs), depending on lines and treatments (Sup. Fig. 8A). In each line, the number of identified TE families was always higher in the WD-RW treatment (Sup. Fig. 8A). Overall, the two different classes of TEs (DNA transposons and retrotransposons) were detected in our mobilome-seq data (Fig. 6A). Most of the eccDNAs identified belonged to the annotated Gypsy, Copia, ENSPM, L1, Ogre, POPGY and SAT superfamilies of TEs and repeats. TEs Depth depth oOf cCoverage (DOC) ranged from 4X to 51000X for the most active TEs, whereas Split Reads (SRs) coverage ranged from 3X to 4600X (Fig. 6A). Reads spanning the two TE extremities constitute an evidence of a circular TE and were detected using a split reads (SRs) mapping strategy. We detected a high number of SRs with a coverage ranged from 3X to 4600X confirming the presence of eccDNAs from TEs (Fig. 6A). The detection of SRs suggested the presence of reads spanning the junction of eccDNAs. Given the range of variation of the DOC coverage, we assigned TE families to four groups following Lanciano et al., (2017). TEs identified in the WT and RNAi lines belonged to the four groups in both WW and WD-RW treatments (Fig. 6A). The “TE^+++^’’group was exclusively represented by the *Gypsy* superfamily, while the ‘‘TE^++^’’group was represented by DNA transposons. In contrast, ‘‘TE^+^’’ and ‘‘TE’’ groups were represented by different superfamilies (Fig. 6A).

**Figure 6:**
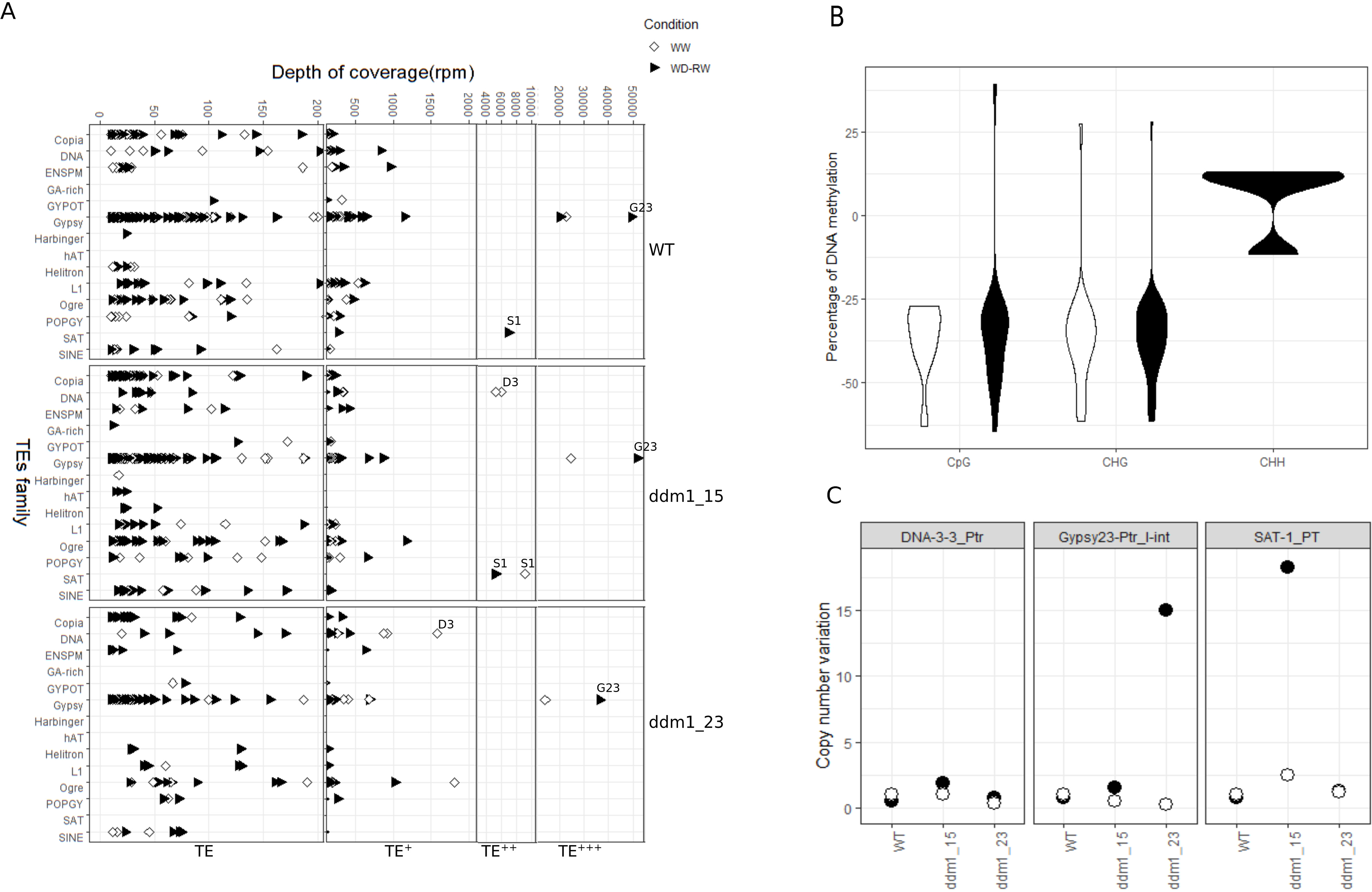
Transposable elements activity among RNAi and WT lines in shoot apical meristem collected one week after rewatering (t2). **A**. Depth of coverage (read per million, rpm) of different TE families in the three different lines and in both WW (white) and WD-RW (black) regimes. TE, TE+, TE++, TE+++ represent different groups of TE families according to their coverage. TE ranges from 0 to 200X; TE+ = 200X – 2000X; TE++ = 2000X - 10000X; TE+++ = 10000X – 55000X. **B**. DNA methylation variations in TEs for each context of methylation in WW (white) and WD-RW (black) conditions. **C**. Copy number variations of three different TEs (DNA-3-3, Gypsy23 and SAT-1) in the three different lines in both WW and WD-RW regimes. White cicles for WW and black circles for WD-RW.

The most active TE (*Gypsy23*, with a DOC ranging from 36000X to 52000X) was present in all lines, but only in the WD-RW treatment (Fig. 6A). The second most active TE also belonged to the *Gypsy* superfamily (*Gypsy27*, DOC ranging from 14000X to 25000X), and was detected in the three lines in the WW treatment but only in the WT line in the WD-RW treatment. We also identified eccDNAs that were line- and/or stress-specific, such as those originating from the satellite *SAT-1* specifically activated in *ddm1-15,* in both WW and WD-RW treatments (9000X and 5200X, respectively), and in the WT line only in the WD-RW condition. Two *DNA-3-3* TEs with the same name, but different in terms of sequence were also detected in the WW treatment only. The first one (named *DNA-3-3_1*) was activated in the two RNAi lines (6000X in *ddm1-15* and 1500X in *ddm1-23*), while the second one (named *DNA-3-3_2*) was specific to *ddm1-15* (5200X) (Fig. 6A). We were also able to detect TEs that belonged to the ‘‘TE^+^’’ group that were line- or stress-specific (Fig. 6A). GO annotation of the genes co-localizing with TEs (+/- 10 kb of genes) revealed enrichment in functions such as response to stress, including hormone response, multicellular organism development, and negative regulation of cellular processes (Sup. Fig. 8B). The number of genes identified in the vicinity of TEs varied between 45 (TEs inside genes) and 1788 (when considering TEs at +/-25kb of genes) (Sup. Fig. 8C). However, only a few of these genes showed changes in their expression level (seven DEGs when considering the vicinity of +/- 25kb; Potri.005G223200, Potri.006G062100, Potri.013G100800, Potri.013G101000, Potri.013G103000, Potri.013G120800, Potri.016G070100).

### Methylation of active TEs and estimation of copy number variations for highly active TEs

About 21% of the mobilome-seq TE families strictly co-localized within common DMRs. This percentage raised to more than 50% when considering the presence of +/- 25 kb of a DMR in the vicinity (Sup. Fig. 8C). These methylated active TE families were in majority hypomethylated (*ca*. 92% of the TE families) in *ddm1* lines (Fig. 6B). In the WW treatment, active TEs were hypomethylated mostly in CG and CHG contexts, while in the WD-RW treatment, active TEs were hypomethylated in CG and CHG contexts but mostly hypermethylated in the CHH context (Fig. 6B). The most active TEs were methylated in CHG context.

To investigate whether TE activity (as detected by eccDNA presence) had led to new integrations in the genome, copy number variation was assessed for three active TEs (*DNA-3-3_1*, *Gypsy23* and *SAT-1*) localized in or near DMRs by qPCR analysis (Fig. 6C). For *DNA-3-3_1*, there was no significant variation in the copy number regardless of the lines and treatments (Fig. 6C). In contrast, an increase in copy number was detected in the WD-RW treatment for *Gypsy23* in *ddm1-23* (15 copies), and for *SAT-1* in *ddm1-15* (18 copies) (Fig. 6C). *Gypsy23* was hypomethylated in the CHG context, while *SAT-1* was found in +/- 2kb of DMRs that were CG and CHG hypomethylated.

## DISCUSSION

### DDM1-dependent DNA methylation plays a role in tree phenotypic plasticity in response to water deficit

#### Hypomethylated poplar RNAi-ddm1 lines are more tolerant to water deficit

Tolerance to water deficit is a complex trait encompassing multiple physiological determinants that can relate to processes as diverse as growth maintenance, survival, or recovery, depending on the context {intensity × duration} considered (McDowell et al., 2008; Volaire et al., 2018). Poplars are among the most sensitive temperate trees to water deficit although significant variation naturally occurs between and within species (Monclus et al., 2006; Street et al., 2006; Fichot et al., 2015). The water deficit imposed in our experiment was deliberately controlled and moderate, as attested by the relatively high Ψ_pd_ (*ca*. -0.8 MPa) and the small effects on net CO_2_ assimilation rates. This avoided rapid growth cessation and instead promoted steady-state acclimation. However, this was sufficient to reveal clear differences in drought responses between the wild type and hypomethylated RNA_i_-*ddm1* lines. While growth was progressively slowed down in the wild type as REW dropped below 40% (Bogeat-Triboulot et al., 2007), both RNAi-*ddm1* lines remained unaffected, suggesting improved tolerance to moderate water deficit. RNA-*ddm1* lines showed a limited increase in time-integrated WUE in response to water deficit as compared to the wild type, which was partly attributable to a less important stomatal closure, at least in *ddm1-15*. The slightly more negative leaf Ψ_min_ observed in RNA_i_-*ddm1* lines tended to confirm the idea that improved tolerance might have been linked partly to a better ability for leaf gas exchange maintenance. In addition, while stomatal density was comparable between the wild type and RNA_i_-*ddm1* lines under standard conditions, stomatal density increased in response to water deficit in the wild type only. *ddm1* mutants in *Arabidopsis* were reported to have more stomata with modified organization (grouped clusters of two or more stomata) (Vassileva et al., 2016), suggesting that *DDM1* modifications have the potential to modify stomatal patterning and therefore leaf physiology. Our findings go further and suggest that these effects may also depend on the environmental context. Interestingly, stomatal density seems to be tightly linked to the dynamics of VPD- and irradiance-induced stomatal closure/opening under water deficit in poplar, with higher densities being correlated with faster stomatal dynamics (Durand et al., 2019). It is therefore possible that the increased stomatal density observed in the wild type under water deficit contributed to a higher stomatal sensitivity, providing one possible explanation for constrained leaf gas exchange and growth reduction as compared to RNA_i_-*ddm1* lines.

Besides differences in dynamic drought responses, our results also suggest that RNA_i_-*ddm1* lines were intrinsically more tolerant to severe water deficit because of higher xylem resistance to drought-induced cavitation. Xylem resistance to drought-induced cavitation is a key trait for plant water relations in a context of survival, as it partly sets operational limits for water transport under tension (Brodribb & Cochard, 2009; Barigah et al., 2013). How modifications of the DDM1 machinery can affect xylem resistance to cavitation remains unknown at this stage. Interestingly, basic xylem properties such as vessel diameter and density, xylem density, or even xylem biochemical composition were not significantly different from the wild type. However, given the mechanistic understanding of drought-induced cavitation in angiosperms (Choat et al., 2018), it is likely that the increased resistance observed in the RNA_i_-*ddm1* lines was primarily linked to modifications of the ultrastructure of vessel-vessel bordered pits (Plavcová Hacke, 2011; Awad et al., 2012; Dusotoit-Coucaud et al., 2014; Herbette et al., 2014). Whether the slight gain in intrinsic cavitation resistance (*i.e.* a few tenth of an MPa) does promote increased survival rates under severe water deficit, and whether epigenetics might be exploited as such for increasing drought tolerance, remains to be purposely tested.

Several DEGs identified in the SAM of RNAi-*ddm1* lines could explain at least in part the improved tolerance to moderate water deficit. Indeed, genes involved in the cuticle and waxes (*CER8*, *FLA12*) that provide protection from abiotic and biotic stresses were upregulated in RNAi-*ddm1* lines. Increased levels of cuticular waxes have been associated with enhanced drought tolerance by preventing uncontrolled water loss (Chen et al., 2011; Wettstein-Knowles, 2016). Similarly, the *MYB106* TF, a major regulator of cuticle formation was upregulated in the *ddm1* line (Oshima and Misuda, 2013, 2016). Other transcription factors (TFs) (*SVP*, *MYB48*, *MYB106, WRKY36, WRKY33, WRKY53*) associated with drought tolerance in plants were also upregulated in the *ddm1* lines (Bechtold et al., 2016; Sun & Yu, 2015; He et al., 2016; Guo et al., 2019). Finally, previous studies in *Arabidopsis ddm1* mutants and derived epiRILs subjected to drought, or more broadly to increased nutrients or salt stress, have provided evidence that variation in DDM1-derived DNA methylation can cause substantial heritable variation in ecologically important plant traits and their plasticity (Reinders et al., 2009; Johannes et al., 2009; Latzel et al., 2013; Cortijo et al., 2014; Zhang et al., 2013, Kooke et al., 2015; Cho et al., 2016; Zhang et al., 2018; Furci et al., 2019).

#### Hypomethylated poplar RNAi-ddm1 lines show leaf symptoms commonly observed in response to pathogen infection

First characterized by Zhu et al., 2013, poplar RNAi-*ddm1* lines did not show any growth or developmental defects when grown under standard greenhouse conditions, but some newly formed leaves displayed a mottled phenotype after a dormancy cycle. This is similar to other reports in *Arabidopsis* and other species, where developmental abnormalities usually appear in later generations (Kakutani, 1997; Tan et al., 2016; Corem et al., 2018; Long et al., 2019). In our RNAi-*ddm1* lines we found two distinct leaf phenotypes: “mottled leaves” and “folded leaves.” However, the two RNAi lines differed in terms of the quantities of symptomatic leaves, with higher values in *ddm1-15* for mottled leaves. This was observed even though our stress treatments had no effect on leaf symptom numbers for either line. The alteration of leaf phenotype has already been attributed to *DDM1* mutation in *Arabidopsis,* where mutants showed altered leaf shape (Kakutani et al., 1995; Qüesta et al., 2013; Kooke et al., 2015), and curled or folded leaves were also observed in hypomethylated *ddc* (*drm1 drm2 cmt3*) mutants (Forgione et al., 2019). However, here we report that stressed conditions do not have an effect on this phenotype.

The presence of mottled leaves could be suggestive of a hypersensitive response (HR) which could result in the hyperactivation of disease resistance genes (Zhu et al., 2013) due to the constitutive hypomethylation in RNAi-*ddm1* lines. This is in agreement with the recent report of Furci et al., (2019) showing that hypomethylation in Arabidopsis *ddm1*-epiRILs at selected pericentromeric regions controls quantitative disease resistance against *Hyaloperonospora arabidopsidis*, which is associated with genome-wide priming of defense-related genes. Lesion mimic mutants, characterized by the formation of necrotic leaves (hereafter called mottled leaves) in the absence of pathogens have been identified, and most importantly, these mutants show enhanced resistance to pathogen infection. This is probably due to mis-regulation of defense responsive genes (Lorrain et al., 2003; Wu et al., 2008). In poplar RNAi-*ddm1* lines, we found an important set of genes involved in defense and immune response that were significantly and differentially expressed in RNAi lines compared to the WT. Immune responsive genes were upregulated in RNAi-*ddm1* lines, while defense related genes were in majority downregulated. Consequently, genes involved in cell death and leaf senescence (*MLO12* and *NHL10*) were upregulated. The activation of immune response genes in RNAi-*ddm1* lines suggests that trees react as if they were attacked by pathogens. This could be particularly important as it could prime *ddm1* lines against future pathogen infections (Lopez Sanchez et al., 2016; Furci et al., 2019). Indeed, different TFs (*WRKY18*, *WRKY33*, *WRKY70*, *WRKY53* and *ERF1*) known to be activated during pathogen attack were overexpressed in *ddm1* lines. The WRKY TFs are known to play important roles in plant defense responses that are related to attacks by several pathogens. Accordingly, overexpression of one of these WRKY TFs in *A. thaliana, Brassica napus* or *Oryza sativa* resulted in enhanced disease resistance (Li et al., 2004; Wang et al., 2014; Wang et al., 2017; Chujo et al., 2007; Marcel et al., 2010; Chen and Chen, 2002; Abeysinghe et al., 2018). In poplar, overexpression of *WRKY18* (with *WRKY35*) also activates pathogenesis-related genes, and increases resistance to the biotrophic pathogen *Melampsora* (Jiang et al., 2017).

The RNAi-*ddm1* lines also had altered expression of a cluster of disease resistance genes, as was already reported in *A. thaliana* (Saze and Kakutani, 2007; Stokes et al., 2002; Lopez Sanchez et al., 2016; Furci et al., 2019). There are multiple mechanisms by which DNA hypomethylation could regulate defense gene induction in *cis* (promoter or nearby TEs) or *trans* contexts. A few *cis*-regulated genes can control the induction of several groups of defense genes by DNA (de)methylation, and hypomethylation can influence chromatin structure at distant genome loci such as TEs activating the RdDM pathway (Grandbastien, 1998; Makarevitch et al., 2015; Quadrana et al., 2019). This recent data in Arabidopsis have shown that methylation controls global defense gene responsiveness *via* trans-acting mechanisms (Lopez Sanchez et al., 2016; Cambiagno et al., 2018; Furci et al., 2019). Accordingly, we found, by overlapping DMRs and DEGs, that only seven DEGs co-localized with DMRs. Expression and methylation values (DEGs at +/-10 kb of DMRs) showed a significant correlation. This suggests that these genes may not be the primary target of *DDM1*.

Active extrachromosomal forms of TEs from several families were identified in water deficit conditions or in *DDM1* lines (mostly hypomethylated). Some of these endogenous TE insertions are located in the vicinity of stress responsive genes (45) and of DEGs (3, at +/-10 kb of TEs). Furci et al., (2019) reported that DNA hypomethylation at TE-rich epiQTLs in *Arabidopsis thaliana* could mediate the induction of defense-related genes across the genome via a *trans*-acting mechanism. Quadrana et al., (2019) further showed that in *Arabidopsis* epiRILs, the insertion of *Copia* elements preferentially targets environmentally responsive genes such as cell death, defense response and immune response, potentially facilitating rapid adaptation. Here, we could detect increased copy numbers for two TE families in the hypomethylated *ddm1* stressed lines. The potential impact of these new insertions on adaptation is unclear. Finally the fact that only two TE families showed increased copy number suggests that additional mechanisms prevent genomic insertions of reactivated TEs.

#### Hypomethylated poplar lines exhibit an altered phytohormonal balance in the shoot apical meristem

Phytohormones are key regulators of plant development and response to stress (Gaillochet & Lohmann, 2015), and have been linked to epigenetic control (Latzel et al., 2012; Yamamuro et al., 2016; Ojolo et al., 2018; Raju et al., 2018), where this interplay could have a major role in meristems for developmental plasticity (Maury et al., 2019). Here, we report that *ddm1* lines exhibited modified shoot apex hormonal balance immediately post-rewatering. Although abscisic acid (ABA) (Zhang et al., 2006; Fernando & Schroeder, 2015) and free auxins (Shi et al., 2014; Basu et al., 2016) are known to mediate physiological responses to water deficit, we found no treatment-induced difference in the SAMs of *ddm1*-lines. In contrast, salicylic acid (SA) and cytokinins (CK) showed significant variations among our *ddm1* lines compared to the WT after drought treatment. Increased SA level was accompanied by a decrease in CK content (especially for zeatine riboside and zeatine-O-glucoside riboside) in post-stress *ddm1* lines. This is in agreement with several reports, which found that drought-stressed plants tend to increase the endogenous level of SA promoting tolerance to several stresses including drought (Munne-Bosch and Penuelas, 2003; Bandurska & Stroi ski, 2005; Azooz & Youssef 2010; Pandey & Chakraborty, 2015; Sedaghat et al., 2017), and decreasing CK levels (Havlovà et al., 2008; Nishiyama et al., 2011; Ha et al., 2012). Altogether our data show that the shoot apex of hypomethylated, stressed lines displayed specific hormonal changes for SA and CK. Shoot apices were collected after three weeks of stress (the day of rewatering) to measure the hormonal balance at the end of the treatment. This timing of sampling could explain why drought-related modifications in ABA or auxin contents were not observed. Correia et al., (2014) have shown in *Eucalyptus globulus* that while drought causes an increase in ABA levels in leaves, rewatering was accompanied by a gradual decrease and stressed and non-stressed plants reached the same amount of ABA seven days post-drought. This also in agreement with Cotrozzi et al., (2017) who showed a very transient peak of ABA following drought conditions in *Quercus ilex*. However, this timing of sampling for hormonal content enabled us to compare it with the presumably (meta)stable epigenetic condition (memory) one week post-rewatering.

DMGs and DEGs nearby active TEs identified in the SAM of *ddm1* stressed lines were mainly involved in development, stress response and phytohormone pathways (such as JA, SA and ethylene). When comparing the genome-wide distribution of DMRs to DEGs in poplar, Lafon-Placette et al., (2018) showed that variations in soil water availability induced changes in DNA methylation preferentially in genes involved in phytohormone metabolism and signaling, potentially promoting phenotypic plasticity. Accordingly, poplar SAMs may also retain an environmental epigenetic memory by targeting hormone-responsive genes (Le Gac et al., 2019). *SAMT1*, a salicylic acid methyltransferase gene was still up-regulated one week post-stress in RNAi-*ddm1* lines compared to the WT, supporting the increase in SA content at the end of the stress. When screening for candidate genes for drought tolerance in *Coffea arabica* cultivars, Mofatto et al., (2016) found that *SAMT1* was upregulated under drought conditions. Zhang et al., (2016) have also shown that *DDM1* affects early seedling growth heterosis in Col/C24 hybrids. Indeed, *ddm1* mutants showed impaired heterosis (Kawanabe et al., 2016) and increased expression of non-additively expressed genes related to salicylic acid metabolism. They proposed that *DDM1* acts as an epigenetic link between salicylic acid metabolism and heterosis, also protecting plants from pathogens and abiotic stress. SA accumulation has also been widely used as a reliable marker of elevated defense responses under pathogen infection, and has been associated with HR cell death or systemic acquired resistance, as well as with *DDM1* mutation (Dong, 2004; Song et al., 2004; Liu et al., 2010; Zhang et al., 2016; Badmi et al., 2019).

The elevated amount of SA levels in RNAi-*ddm1* lines could be suggestive of acquired disease resistance in *ddm1* mutants. Similarly, two genes (*CYP94B1* and *OPR2*) involved in JA metabolism were upregulated in the RNAi-*ddm1* lines. *CYP94B1* and *OPR2* have been shown to contribute to the attenuation of the JA-dependent wound responses in response to pathogen attack (Koo et al., 2014; Pandey et al., 2017; Wasternack & Hausse, 2018). Moreover, Latzel et al., (2012) have established a positive correlation between DNA methylation variations (epiRILs) and JA and SA responses, and proposed that part of the variation of plant defenses observed in natural populations may be due to underlying epigenetic, rather than entirely genetic, variation. Several TFs involved in abiotic (drought) or biotic stress, and acting in phytohormone pathways, were upregulated in stressed *ddm1* lines in comparison to WT; this includes genes such as *SVP* (*SHORT VEGETATIVE PHASE*) that can confer drought resistance by regulating ABA catabolism (Wang et al., 2018), *MYB48* that can improve tolerance to drought when overexpressed (Wang et al., 2017) and *ERF1,* which can activate a subset of ethylene-inducible genes as the immune responsive gene *PDF1* (Solano et al., 1998; Fujimoto et al., 2000; Heyman et al., 2018).

Altogether our data suggest a direct connection between epigenetic regulation and phytohormones in the meristem for the control of plasticity as previously proposed (Maury et al., 2019). This interplay could control the expression of cell identity genes, the stable activation of hormone-responsive genes post-stress, or act as an integrative hub for the sensing of hormonal balance to ensure plasticity, and potentially environmental memory (Maury et al., 2019).

### DDM1-dependent DNA methylation in poplar shoot apical meristem is context dependent and affects both genes and TE activity

#### DDM1-dependent DNA methylation is cytosine context dependent

Poplar *ddm1* RNAi knock down lines were hypomethylated in SAMs according to global DNA methylation levels (7 to 17% reduction compared to WT). This is in agreement with both the WGBS analysis, which showed DMRs mostly hypomethylated, and with the report of Zhu et al., (2013), who found comparable reductions in leaves. While poplar *DDM1*-dependent DNA methylation was affected in the three contexts (CpG, CHG & CHH), strong differences were observed among the contexts. The methylation level in CHG was drastically reduced compared to CpG or CHH contexts, suggesting that in poplar, *DDM1* preferentially targets methylation in the CHG context. This is in accordance with rice (Tan et al., 2016), and maize (Li et al., 2014; Long et al., 2019), but contrasts with Arabidopsis and tomato, where the disruption of *DDM1* led to a drastic hypomethylation of the genome, mainly in CpG and CHG contexts (Vongs et al., 1993; Kakutani et al., 1995, 1997; Lippman et al., 2004; Zemach et al., 2013; Corem et al., 2018). These observations suggest that *DDM1* has differential effect on DNA methylation patterns in diverse species (Tan et al., 2016; Long et al., 2019).

We also found that *DDM1* reduction caused extensive CpG, and to a lower extent CHG, hypermethylation of intergenic regions and genes, and that CHH hypermethylation of TEs only occurred in stressed conditions. The loss of *DDM1* has already been associated with hypermethylation in CHG contexts in genes of Arabidopsis and rice, as well as more limited hypermethylation in both CpG and CHG contexts for euchromatic regions in tomato and maize (Lippman et al., 2004; Mathieu et al., 2006; Saze & Kakutani, 2007; Zemach et al., 2013; Tan et al., 2016; Corem et al., 2018; Long et al., 2019). CHH hypermethylation has also been reported for heterochromatic TEs in rice and tomato (Tan et al., 2016; Corem et al., 2018). Here, we found that extensive CpG hypermethylation, notably in genes, was not associated with differential expression level, and overlapped with only a few active TEs, while hypermethylated CHH TEs showed reduced activity in comparison to hypomethylated CG and CHG TEs. It has been proposed that these hypermethylation events are likely to be mediated by different, potentially overlapping mechanisms that act as an internal balancing mechanism to compensate for the extensive loss of methylation in other contexts (Zemach et al., 2013; Tan et al., 2016; Corem et al., 2018).

Altogether, we propose that hypermethylation events, being species-dependent, may participate in pleiotropic effects. This could be related to variations in the functional role of *DDM1* in the different species, and/or in relation to their epigenetic machinery and genome complexity. We provide evidence that this phenomenon is also stress-dependent, and occurs in the shoot apical meristem with possible implications for mitotic and meiotic transmission.

#### DDM1 impairment of gene methylation has limited effects on gene expression

*DDM1* decrease in poplar affected the methylation of 879 and 910 genes in WW and WD-RW treatments, respectively. In poplar, gene methylation typically occurs in CpG and CHG contexts, and to a lower extent in the CHH context (Feng et al., 2010; Vining et al., 2012; Lafon-Placette et al., 2013). Despite a strong reduction in genome-wide CHG methylation, the numbers of DMGs found in CpG and CHG were of similar order. However, in CHG and CHH contexts genes found were in majority hypomethylated (>90%), whereas in the CpG context an equal number of genes were hypo- or hypermethylated.

Transcriptomic analysis in SAMs only revealed a limited number of DEGs (136) between the WT and *ddm1-23* in response to water deficit, with 76 upregulated and 60 downregulated genes. Previous data on Arabidopsis and tomato *ddm1* mutants were of similar magnitude (Zemach et al., 2013; Corem et al., 2018), while more DEGs were reported in rice (Tan et al., 2016) or in infected Arabidopsis EpiRILs (Furci et al., 2019). Among DEGs, only seven strictly overlapped with DMRs (up to 39% for genes at ∼ +/-10 kb of DMRs); in agreement with tomato *ddm1* mutants (Corem et al., 2018). This suggests that not all of these DEGs are primary targets of DDM1. A significant negative rank correlation was detected, however, between gene expression and methylation for the 53 DEGs located at less than +/-10 kb of DMRs. Although changes in 5’ DNA methylation may influence gene expression (Seymour & Becker, 2017), the role of gene- body methylation still remains disputed (Bewick & Schmitz, 2017), and to date it has been difficult to differentiate between direct changes mediated by DNA methylation and secondary effects (Meyer, 2015). In most available studies, no correlation could be detected between DNA methylation and expression changes at the genomic level. However, in agreement with other studies, our data show that the transcriptional activity of a subset of genes might be regulated by DNA methylation in response to abiotic stress (Karan et al., 2012; Garg et al., 2015; Chwialkowska et al., 2016, Lafon-Placette et al., 2018). These genes, including TFs and hormones-related pathways, are likely to explain, at least in part, the developmental plasticity of poplar *ddm1* lines (Maury et al., 2019).

#### DDM1-dependent DNA methylation favors TEs reactivation and insertion during stress

The role of DNA methylation and DDM1 on TE proliferation is well-established in plants (Miura et al., 2001; Mirouze et al., 2009; Tsukahara et al., 2009). TE proliferation has also been investigated in DDM1-epiRILs in Arabidopsis and tomato (Reinders et al., 2009; Johannes et al., 2009; Corem et al., 2018; Quadrana et al., 2019). Here, we reported TE activity in the SAM of WT and RNAi-*ddm1* lines using the mobilome-seq workflow (Lanciano et al., 2017). TEs, which represent ∼42 % of the genome of poplar (Kejnovski et al., 2012), were evaluated both under control and post water deficit conditions. The pattern of TE methylation varied widely between the different contexts. While most TEs were hypomethylated in CG and CHG contexts, we could not detect any TEs overlapping with the common DMRs, suggesting a limited effect in the CHH context (Corem et al., 2018). This could be related also to the threshold that we applied during DMR identification, which considered only DMRs with at least 10% of difference and 10X of coverage. This is very stringent since CHH methylation level in poplar is very low compared to CG and CHH contexts (∼3,25%; Feng et al., 2010). This result also supports a redundant function of CG and non-CG methylation in the transcriptional silencing of the TEs (Ikeda & Nishimura, 2015). The burst of TEs in *ddm1* lines were recorded for the two different classes (DNA transposons and retrotransposons), with a notable enrichment for the retrotransposons of the *Gypsy* family. Interestingly, when assessing the copy number variation of the most active TEs, we could detect only increased copy number for *Gypsy* retrotransposons (*Gypsy-23* and *SAT-1/Gypsy-78_Ptr-I-int*) during the post stress episode, suggesting specific control of *Gypsy* activity by DDM1 during the stress. *Gypsy* elements are long terminal repeat (LTR)-flanked retrotransposons that are concentrated in pericentromeric heterochromatin, in comparison to other repeats that are more dispersed e,g,m (*Copia*, *LINE*, *SINE*). Wang et al., (2018) recently reported a constant conflict between *Gypsy* retrotransposons and CHH methylation within a stress-adapted mangrove genome, and found differential accumulation among classes of LTR TEs mainly due to siRNA-mediated CHH methylation preferentially targeting *Gypsy* elements. This is consistent with our data as we observed extensive CHH methylation of TEs (such as *Gypsy* elements) during drought in the *ddm1* lines associated with enrichment in non-coding RNA (ncRNA) GO labels. This may suggest that *ddm1* lines tend to methylate TEs in CHH contexts in order to repress their activity. Wang et al., (2018) subsequently proposed that the apparent conflict between TEs activity and repression of integration may enable the maintenance of genetic diversity and thus evolutionary potential during stress adaptation. This is particularly interesting as genes found with nearby active TEs in *ddm1* lines are mainly involved in stress response and development. Quadrana et al., (2019) recently proposed that TEs are potent and episodic (epi)mutagens that increase the potential for rapid adaptation. They proposed that this is in large part due to epigenetic mechanisms of suppression that limit their activity which might result in purifying selection against them, and also limits their mutation rate due to their presence in highly compact chromatin. Here our data suggest that DDM1 could play a role in these repressive but diversity maintaining mechanisms during drought stress in poplar.

### Shoot apical meristem is a central place for epigenetic control of developmental plasticity

Our study focused on meristems, previously described as “organs with specific epigenetic machinery” (Baubec et al., 2014; Lafon-Placette et al., 2018; Le Gac et al., 2018, 2019) and as controlling centers of development and acclimation. They are also the loci of mitotic and meiotic transmission. Here, we report the detailed characterization from physiological to omics levels of two independent *ddm1* poplar RNAi lines. Both lines exhibited higher tolerance to drought, mottled leaves, and modification of hormonal balance. Our study shows that *DDM1*-dependent DNA methylation in the shoot apical meristem of poplar trees plays two roles: controlling developmental plasticity and enabling stress response through a direct interplay with hormonal pathways (Maury et al., 2019). *DDM1*-dependent DNA methylation also controls the activation, and probably the integration, of TEs whose movements can induce heritable mutations and affect the potential for rapid adaptation (Kawakatsu & Ecker, 2019; Quadrana et al., 2019). These results are in agreement with previous studies in poplars (Gourcilleau et al., 2010; Zhu et al., 2013; Conde et al., 2017; Lafon-Placette et al., 2018; Le Gac et al., 2018, 2019; Sow et al., 2018) and recent findings in annuals (Raju et al., 2018; Schmid et al., 2018; Zhang et al., 2018; Furci et al., 2019; Quadrana et al., 2019), both showing that epigenetic variation and TEs have the potential to create phenotypic variation that is substantial, persistent, and stable, thus of adaptive and evolutionary significance.

Our data are consistent with recent models about role of epigenetic variation in plants (Yona et al., 2015; Richards et al., 2017; Kawakatsu & Ecker, 2019; Maury et al., 2019) that view meristems as an interface between physiological response and genetic adaptation. Further studies are needed to examine the role of stress-induced epigenetic variation and associated mutation that examine a far greater range of stresses, species, and genotypes. This will enable its importance relative to more traditional sources of adaptive and evolutionary variation to be interpreted. Such studies are of particular interest for long-lived organisms in the age of rapid, anthropogenic climate change.

## METHODS

### Plant material, experimental design, and control of water deficit

Experiments were conducted on two *PtDDM1* RNAi lines (*ddm1-*15 and *ddm1-*23), and a wild type (WT) line of *Populus tremula × Populus alba* (clone INRA 717-1B4). These two RNAi lines were chosen among those previously described by Zhu et al., (2013) for consistently lower levels of both cytosine methylation (17.0 and 16.7% reduction compared to WT, respectively) and *PtDDM1* residual expression (*ca.* 62% reduction). Six week-old *in vitro* propagated plantlets were first transferred into small sealed chambers for progressive acclimation. Acclimated plantlets were then transferred to 4L pots filled with a potting substrate (Klasmann RHP 25-564, pH = 5.8) complemented with Osmocote PG Mix (1 kg/m^3^ of N-P-K 80/35/60). The experiment was conducted in a greenhouse located at the research station of INRAE Orléans Centre Val-de-Loire (47°46’N, 1°52’E), with a photoperiod of 16:8, an average temperature of 21°C, and a relative humidity of 32%. Right before the water deficit experiment started (2 month-old plants), three plants per line were randomly sampled (t_0_). The remaining 81 plants were then randomly distributed into nine blocks (*n* = 3 trees per line per block) and assigned to either a well-watered control treatment (WW, n = 1 per line per block) or a water deficit treatment followed by re-watering (WD-RW, n = 2 per line per block).

Water deficit was initiated at t_0_ and lasted three weeks until t_1_. Watering was performed every two days and was adjusted for each plant based on volumetric soil water content (SWC) estimated *via* pot weighing. Values of SWC at time i (SWC_i_) were converted to soil relative extractable water (REW_i_, %) using the following equation: REW_i_ = (SWC_i_ – SWC_wp_) / (SWC_fc_ – SWC_wp_) × 100 where SWC_fc_ and SWC_wp_ correspond to the SWC at field capacity and at the wilting point, respectively. Plants from the WW treatment were always watered to field capacity, while plants from the WD-RW treatment were re-watered to a target value of approx. 40% of REW. At t_1_, plants from the WD-RW treatment were re-watered to field capacity, three blocks were sampled, and the remaining six blocks were maintained watered for one week until t_2_, after which all remaining plants were sampled. The water deficit intensity was evaluated at t_1_ by measuring the predawn leaf water potential (Ψ_pd_, MPa) during the night preceding the re-watering. Measurements were performed on a subset of five randomly selected blocks using a pressure chamber (PMS instruments, Albany, OR, USA). Minimum leaf water potential (Ψ_min_) was estimated for the same plants at midday on the day preceding re-watering.

### Physiological and phenotypic characterization

#### Growth and leaf symptoms

Stem height was measured every two days using a telescopic ruler, while stem diameter was measured every four days using a digital caliper. Due to Zhu et al. (2013) reporting the occurrence of spontaneous necrotic spots on the leaves of *ddm1* RNAi lines (mottled phenotype), we repeatedly measured the number of leaves showing necrotic symptoms (mottled leaves) during the whole duration of the experiment. We also counted the number of leaves showing a ‘folded’ morphology (see results section). These measurements were performed on three randomly selected blocks.

### Leaf gas exchange, bulk leaf carbon isotope composition (**δ**^13^C), and stomatal density

Leaf gas exchange were measured using a LI-6400 open path photosynthesis system (Li-Cor, Lincoln, NE, USA) equipped with an LED light source (LI-6400-02B). Measurements were systematically performed on fully mature leaves in the top third of the plant in the same five blocks as those used for Ψ_pd_ and Ψ_min_. From the time of drought initiation onwards, net CO_2_ assimilation rate (A_net_, µmol m^-2^ s^-1^), and stomatal conductance to water vapor (g_s_, mol m^-2^ s^-1^) were measured every day between 9 am and 3 pm in order to characterize the dynamic response of the genotypes to water deficit. Measurements were performed at a saturating photosynthetic photon flux density (PPFD) of 2000 µmol m^-2^ s^-1^, an ambient CO_2_ concentration of 400 ppm, a constant block temperature of 25°C, and a reference vapor pressure deficit (VPD) maintained close to 1kPa.

Bulk leaf carbon isotope composition (δ^13^C) was used as a time-integrated surrogate of leaf intrinsic water-use efficiency (Farquhar et al., 1982). Six calibrated discs (3.14 cm^2^) of leaf lamina were punched from a mature leaf at t_2_ on all plants. Leaf disks were oven-dried at 60°C for 48hrs before being ground to a fine powder. One milligram subsamples were then enclosed in tin capsules and combusted at 1200°C. The CO_2_ produced by combustion was purified, and its ^13^CO_2_/^12^CO_2_ ratio was analyzed using isotope ratio mass spectrometry (IRMS, Finnigan MAT Delta S, Bremen, Germany). The δ^13^C (‰) was expressed relative to the Vienna Pee Dee Belemnite standard and calculated as δ^13^C = (R_sa_-R_sd_)/R_sd_ × 1000 where R_sa_ and R_sd_ are the ^13^CO_2_/^12^CO_2_ ratios of the sample and the standard, respectively (Farquhar et al., 1989). The accuracy of the δ^13^C measurements done by IRMS during the time samples was assessed using referenced standards of ± 0.05 ‰ (SE). All measurements were performed at the INRAE-Nancy technical platform of functional ecology in France.

Stomatal counts were measured on a subset of three blocks, following the method described by Xu & Zhou (2008). Stomatal imprints were taken between the central leaf vein and the leaf edge on both adaxial and abaxial sides, and then fixed on microscopic slides using scotch tape. Slides were observed under a light microscope coupled to a Moticam 580 5.0MP digital camera, and three pictures were taken on each filmstrip side.

### Xylem structure, function and biochemical composition

Xylem vulnerability to drought-induced cavitation was assessed at t_2_ on the well-watered plants of all blocks (INRAE Phenobois Platform, Clermont-Ferrand, France). Vulnerability to cavitation sets the operational limit of xylem under drought, and is a key trait involved in drought tolerance (Brodribb & Cochard, 2009). We used the Cavitron technique which is well suited to poplars (Cochard et al., 2005, Fichot et al., 2015). In short, the technique uses the centrifugal force to increase xylem tension (Ψ_x_, MPa) in stem segments, while at the same time measuring the percent loss of hydraulic conductance (PLC, %). The dependence of PLC upon Ψ_x_ was used to generate vulnerability curves for each stem segment. The following sigmoid function was fitted to data (Cochard et al., 2007): PLC = 100 / (1 + exp((s/25)×(P-P_50_)), where P_50_ is the xylem tension causing 50% loss of hydraulic conductance (MPa) and s is the slope of the curve at the inflexion point (%. MPa^-1^). Values of P_50_ were used as proxies for vulnerability to xylem drought-induced cavitation.

Xylem histology was performed on stem segments of all plants at t_2_. Stem cross-sections 30 µm-thick were obtained using a hand microtome (RM 2155, Leica Microsystems, Vienna, Austria), and stained with safranin (1% in 50% ethanol), and followed by Astra blue (1% in 100% ethanol), before being permanently mounted in Canada Balsam. Stained sections were examined under a light microscope coupled to a Moticam 580 5.0MP digital camera. Pictures covering pith to cambium were taken at a 10x magnification on three opposite radial sectors in order to estimate vessel diameter (µm), vessel density (mm^-2^), vessel lumen fraction (%), and theoretical xylem specific hydraulic conductivity (Fichot et al., 2010). Image analyses were all performed using the ImageJ software (Schneider et al., 2012). Xylem density was assessed on the same stem segments using 4 cm-long samples. Measurements were realized using the Archimedes principle. Bark-free stems were split longitudinally in order to remove wood pith, submerged to estimate the volume of displaced water, and weighed after being oven-dried at 105°C. Xylem density (g.cm^-3^) was computed as the ratio between dry mass and the volume of displaced water (Fichot et al., 2010).

Xylem biochemical composition was evaluated indirectly on stem powders of all plants by using Fourier-Transform mid-Infrared Spectroscopy (FTIR, Spectrum 400, Perkin Elmer, Massachusetts, USA), and home-made calibration models previously developed at the INRAE Genobois phenotyping platform for Klason lignin content, S/G ratio and tension wood content (*unpublished data*). For each sample, a few milligrams were placed three times on the attenuated total reflectance (ATR) diamond for the scan. Spectra ranged from 650 to 4000 cm^-1^ wave numbers with a step of 2 cm^-1^. Spectrum analyses were realized using the R software (R Core Team, 2015). Spectra were first cut and smoothed by using the ‘prospectr’ package before applying normalization and first derivative processing (Bertrand & Dufour, 2006).

### Phytohormone quantification

Shoot apices were immediately frozen in liquid nitrogen upon sampling, and later ground to a fine powder in an automatic ball mill (MM 200 Retsch, Germany). Phytohormone assays for abscisic acid (ABA), free auxin, salicylic acid (SA), jasmonic acid (JA) and cytokinins were performed on the SAMs collected at t1, using LC-MS according to a published procedure (OVCM platform, IJPB, INRAE Versailles, France; Li-Marchetti et al., 2015; Trapet et al., 2016). For each sample, 10 mg of dry powder was extracted with 0.8 mL of acetone/water/acetic acid (80/19/1 v:v:v). Phytohormone stable labelled isotopes used as internal standards were prepared as described in Roux et al., (2014). Two ng of each and 0,5 ng of cytokinines standard was added to the sample. The extract was vigorously shaken for 1 min, sonicated for 1 min at 25 Hz, shaken for 10 minutes at 10°C in a Thermomixer (Eppendorf®), and then centrifuged (8000 g, 10 °C, 10 min.). The supernatants were collected, and the pellets were re-extracted twice with 0.4 mL of the same extraction solution, then vigorously shaken (1 min), and sonicated (1 min; 25 Hz). After the centrifugations, the three supernatants were pooled and dried (Final Volume 1.6 mL). Each dry extract was dissolved in 100 µL of acetonitrile/water (50/50 v/v), filtered, and analyzed using a Waters Acquity ultra performance liquid chromatograph coupled to a Waters Xevo Triple quadrupole mass spectrometer TQS (UPLC-ESI-MS/MS). The compounds were separated on a reverse-phase column (Uptisphere C18 UP3HDO, 100*2.1 mm*3µm particle size; Interchim, France) using a flow rate of 0.4 ml min^-1^, and a binary gradient: (A) acetic acid 0.1% in water (v/v) and (B) acetonitrile with 0.1% acetic acid, and a column temperature of 40°C. Mass spectrometry was conducted using electrospray, and Multiple Reaction Monitoring scanning mode (MRM mode), in either positive ion mode (for the indole-3-acetic acid and cytokinins) or negative ion mode (for the other hormones). Relevant instrumental parameters were set as follows: capillary 1.5 kV (negative mode), with source block and desolvation gas temperatures at 130 °C and 500 °C, respectively. Nitrogen was used to assist the cone and desolvation (150 L.h^-1^ and 800 L.h^-1^, respectively), argon was used as the collision gas at a flow of 0.18 ml.min^-1^.

### DNA extraction and determination of global DNA methylation levels by HPLC

Genomic DNA was extracted from all SAMs with a CTAB protocol (Doyle & Doyle, 1987), and stored at −80°C. Quantity and quality were approximated using a NanoDrop spectrometer (NanoDrop Instrument, France).

For the determination of global DNA methylation, genomic DNA was enzymatically hydrolyzed into nucleosides, and analyzed by high-performance liquid chromatography (HPLC), as described by Zhu et al., (2013). Controls for this procedure included co-migration with commercial standards (Sigma-Aldrich), confirmation by enzyme restriction analysis, and tests for RNA contamination based on the HPLC detection of ribonucleosides. Global DNA methyl cytosine percentages (% mC) were estimated as follows: %mC = (mC/(C + mC)) × 100, where ‘C’ is 2’-déoxycytidine content, and ‘mC’ is 5-methyl-2’-déoxycytidine content. For each line in each treatment, three biological replicates were randomly chosen out of the six for analyses, with two independent genomic DNA extractions per replicate, three hydrolysis replicates and two HPLC runs.

### Whole Genome Bisulfite Sequencing (WGBS) and bioinformatic pipeline for methylome characterization

An equimolar pool of 2 µg DNA at approximately 100 ng/µl, and extracted from four SAMs was made for each line in each treatment. Whole-genome bisulfite sequencing was performed by the CNRGH laboratory (J. Tost, Evry, France) in accordance with the published procedure (http://www.nugen.com/products/ovation-ultralow-methyl-seq-library-systems) adapted from Daviaud et al., (2018). The workflow of the library preparation protocol follows the classical library preparation protocol in which methylated adaptors are ligated to the fragmented DNA prior to bisulfite conversion. A total of 200 ng of genomic DNA was fragmented to a size of approximately 200 base pairs (bp), and then purified and methylated adaptors compatible with sequencing on an Illumina HiSeq instrument were ligated. The resulting DNA library was purified and bisulfite converted. A qPCR assay was used to determine the optimal number of PCR amplification cycles (between 10 and 15 cycles) required to obtain a high diversity library with minimal duplicated reads prior to final library amplification. The sequencing was performed with paired ends (2×150bp) on an Illumina HiSeq4000 platform. Raw data were stored in FASTQ files with a minimal theoretical coverage of 30X (SRA record is under the reference PRJNA611484; https://www.ncbi.nlm.nih.gov/sra/PRJNA611484).

The bioinformatics pipeline used in this study is adapted from the ENCODE pipeline (https://www.encodeproject.org/wgbs/) and installed on the Galaxy instance, accessible of IHPE (http://galaxy.univ-perp.fr/, Perpignan, France). First, quality control and cleaning of the raw data were carried out by only considering nucleotides with a quality score over 26, and reads which had more than 95% of their nucleotides over this quality threshold. The second step was the alignment of the WGBS reads on the reference genome *Populus tremula × alba* (http://aspendb.uga.edu/index.php/databases/spta-717-genome), by using BISMARK (version 0.16.3, Krueger & Andrews, 2011) and bowtie 2 tools (version 2.1.0; Langmead et al., 2009) or a bisulfite sequence mapping program (BSMAP version 2.74) (Xi & Li, 2009). The parameters were modified for ‘paired-end’ alignment, read lengths were between 70 and 500 bp, and others were kept by default (See sup. Figure 1A, B and C). The methylkit R package allowed the identification of DMRs among lines and treatments. A DMR was defined as a 500 bp region with a minimum coverage of 10X, which highlighted the differential methylation between two samples of at least 10% for CHH, and 25% for CHG and CG (q-value=0.01). Different types of DMRs were identified and are referred to as follows: stress-specific DMRs refer to DMRs between WW and WD-RW treatments, line-specific DMRs refer to DMRs between *ddm1* lines and the WT, and common DMRs refer to line-specific DMRs common to both lines *ddm1-15* and *ddm1-23*.

DMR annotation was realized by using reference data available for the *Populus tremula × alba* genome from the aspendb database. Gene Ontology (GO) term enrichment was assessed for the methylated genes with Revigo (http://revigo.irb.hr/) software, using default parameters. ‘TreeMap’ view was performed with rectangle size adjusted to reflect the absolute log10 *P*-value of the GO term, through use of the corresponding poplar model genes from the best v3.0 blast hits with *Arabidopsis* TAIR10 annotations.

### Transcriptomics and bioinformatic pipeline

Total RNA (three biological replicates) was extracted from the wild type and one RNAi line in WD-RW condition by using a modified protocol of Chang et al., 1993). *Ddm1-23* was chosen as the most representative of the two lines as it exhibited a lower decrease in methylation compared to *ddm15-7,* but nonetheless most of its DMRs were in common among the two lines. In brief, SAMs were ground into fine powder in liquid nitrogen and total RNAs were extracted using a CTAB buffer (Changet al., 1993). RNA was precipitated using lithium chloride (10M) and purified using the Macherey Nagel Nucleospin RNA kit (740955). Sequencing was done using the Illumina NexSeq500 (IPS2 POPS platform, Saclay, France). RNA-seq libraries were performed by following the TruSeq_Stranded_mRNA_SamplePrep_Guide_15031047_D protocol (Illumina®, California, USA). The RNA-seq samples have been sequenced in paired-end (PE) with a sizing of 260 bp and a read length of 75 bases. 6 samples by lane of NextSeq500 using individual bar-coded adapters and giving approximately 15 millions of PE reads by sample are generated. To facilitate comparisons, each sample followed the same steps from trimming to counts. RNA-Seq preprocessing included trimming library adapters and performing quality controls. The raw data (fastq) were trimmed with the Trimmomatic (Bolger et al., 2014) tool for Phred Quality Score Qscore >20, read length >30 bases, and ribosome sequences were removed with the sortMeRNA tool (Kopylova et al., 2012). The genomic mapper STAR (version 2.6, Dobin A. et al 2013) was used to align reads against the *Populus tremula × alba* hybrid genome (genotype INRA 717-1B4), with the options outSAMprimaryFlag AllBestScore--outFilterMultimapScoreRange 0, to keep only the best results. The abundance of each gene was calculated with STAR, with paired-end reads only being counted where the reads unambiguously mapped one gene, and multi-hits were removed. Following this pipeline, around 92% of PE reads were associated to a gene, 3 to 4% PE reads were unmapped and 3 to 4% of PE reads with multi-hits were removed.

Differential analyses followed the procedure described in Rigaill et al., (2016). In brief, genes with less than one read, after a count per million normalization in at least one half of the samples, were discarded. Library size was normalized using the trimmed mean of M-value (TMM) method, and count distribution was modeled with a negative binomial generalized linear model. Dispersion was estimated by the edgeR method (Version 1.12.0, McCarthy et al., 2012) in the statistical software ‘R’ (Version 3.2.5 R Development Core Team (2005)). Expression differences compared two samples using the likelihood ratio test, and p-values were adjusted by the Benjamini-Hochberg procedure to control False Discovery Rate (FDR). A gene was declared differentially expressed if its adjusted p-value was lower than to 0.05.

All steps of the experiment, from growth conditions to bioinformatic analyses, were managed in CATdb database (Gagnot et al., 2008; http://tools.ips2.u-psud.fr/CATdb/) with ProjectID NGS2017-01-DDM1 This project is submitted from CATdb into the international repository GEO (Gene Expression Omnibus, Edgard R. et al. 2002, http://www.ncbi.nlm.nih.gov/geo) with ProjetID GSE135313.

### Mobilome-seq and copy number variation of TEs

According to their mode of transposition, TEs generate extrachromosomal circular DNAs (eccDNAs) when active. The sequencing of these eccDNAs by the mobilome sequencing was a successfully method to identify active TEs (Lanciano et al., 2017). Inorder to identify the effect of DDM1 knock down and a hydric stress on the release of TE activity we used approximately 6 µg of genomic DNA to perform mobilome-seq libraries. eccDNAs were isolated and libraries were prepared and sequenced following Lanciano et al., (2017). Bioinformatics was carried out on the *Populus tremula × alba* genome (SPta717 v1.1) by using the same pipelines as described in Lanciano et al., (2017). In order to obtain TE database for *Populus tremula × alba* genome, we used the TE database based on *Populus trichocarpa* genome (version 3.0). In brief, the sequencing reads were first filtered against the mitochondria and chloroplast genomes before being i) mapped against the *P. tremula × alba* reference genome using Bowtie2, ii) mapped for split reads (SR) using segemehl software (Hoffmann et al., 2014) and iii) *de novo* assembled using a5-miseq (Coil et al., 2015). Given the range of variation of the DOC coverage, we assigned TE families to four groups following Lanciano et al., (2017): The first group was named “moderate or not active TEs (group “TE”)”, and comprised TEs with a DOC ranging from 4 to 199X. The second group was named “active TEs” (group “TE^+”^) and comprised TEs with a DOC ranging from 200-1999X. The third group wasnamed “very active TEs” (“TE^++”^) and comprised TEs with a DOC ranging from 2000 to 9999X. The fourth group was named “highly active group” (“TE^+++”^), and comprised TEs with a DOC ranging from 10000 to 51000X. Raw and processed data are available with the GEO accession number GSE147934 (https://www.ncbi.nlm.nih.gov/geo/query/acc.cgi?acc=GSE147934).

Copy number variation of TEs was assessed for all studied lines in both treatments by qPCR using genomic DNA extracted from SAMs. In summary, primers of TEs were first designed using corresponding FASTA sequences with Eprimer3 and Netprimer for quality control checking (http://www.bioinformatics.nl/cgi-bin/emboss/eprimer3, http://www.premierbiosoft.com/NetPrimer). Primers that passed quality control (Gypsy: F=AAC-AAG-CTG-AAG-CCC-AAG-AA, R=TCG-ACC-TCG-AGT-TAG-GTT-CC; DNA-3-3: F=TAG-TGT-GCA-GTG-GAG-CAT-GG, R=AAA-AGC-AGG-GTG-TTT-TGC-TG; SAT-1: F=TCA-CCG-GAA-CCC-ACT-TCT-AC, R=GCA-ACG-ACT-GAG-TTT-CGT-CA) were used with 10 ng/µl of genomic DNA for qPCR analyses (Platinum™ SYBR™ Green qPCR SuperMix-UDG, Invitrogen™ kit). A standard cycling program was applied (50 °C for 2 min, 95 °C for 2 min, and 40 cycles of 95 °C for 15 sec and 60 °C for 30 sec). Melting curves were obtained using recommended qPCR instrument settings. Copy number variation was assessed by using absolute quantification of cycle threshold values (Schmittgen & Livak, 2008).

### Statistical analyses

Statistical analyses were performed with R statistical software under R Studio integrated development environment (R Core Team, 2015, RStudio: Integrated Development for R. RStudio, Inc., Boston, MA URL http://www.rstudio.com/). Means are expressed with their standard errors (SE). Differences between lines and treatments for phenotypic traits were evaluated by analysis of variance (ANOVA) on individual values adjusted for block effects. Tukey’s post-hoc test was used to identify differences between groups when ANOVAs indicated significant effects. Statistical tests were considered significant at *P* < 0.05.

## Author contributions

S.M designed and coordinated the research. The plant experimental design was established by S.M, S.S, R.F, and F.B. Ecophysiological measurements were performed by A.L.L.G, R.F, A.D, I.L.J and H.C; analysis was realized by A.L.L.G and R.F. NIRS measurements and analysis were performed by A.L.L.G and V.S. Phytohormones analysis was performed by S.C. DNA, RNA extractions were done by A.D and A.L.L.G and M.D.S. HPLC analysis was done by A.D and S.M. RNA-seq was realized and analyzed by J.C, V.B, L.S.T with M.D.S. J.T, C.D realized WGBS analysis. WGBS data analysis was done by S.M, A.L.L.G and M.D.S. Bioinformatics for WGBS was done with the help of C.C and C.G. Mobilome analysis was done under the supervision of M.M with S.L and A.L.L.G. QPCR analysis were done by M.C.L.D with A.D and M.D.S. Data analysis was done by S.M, A.L.L.G and M.D.S. Statistical analyses were done by R.F, A.L.L.G and M.D.S. S.M, R.F, A.L.L.G and M.D.S conceived and wrote the first draft of the manuscript. All authors approved the final version of the manuscript.

## Acknowledgments

M.D.S and A.L.L.G received Phd grants from the Ministère de la Recherche et Enseignement Supérieur. This work was supported by INRAE (grant PI EFPA-2014 to S.M.), the RTP3E CNRS grants for mobility (http://rtp-3e.wixsite.com/rt3e; A.L.L.G, S.M, C.G and M.M) and access to the IHPE platform (http://galaxy.univ-perp.fr/, Perpignan, France). The LBLGC also benefits from support of the ANR EPITREE (ANR-17-CE32-0009-01) to SM. English editing was done by Infinity English (N° SIRET 82950710200012). The IJPB benefits from the support of Saclay Plant Sciences-SPS (ANR-17-EUR-0007). This work has benefited from the support of IJPB’s Plant Observatory technological platforms. SL and MM were supported by the French National Agency for Research (ANR-13-JSV6-0002 “ExtraChrom”) and the Laboratoire d’Excellence LABEX TULIP (ANR-10-LABX-41).

## Supplemental materials

**Supplemental Figure 1:** Strategies for methylome bioinformatic analysis **A**. Impact of different types of quality controls on the percentage of mapped reads on two distinct genomes: *P. trichocarpa* and *P. tremula × P. alba.* Dots shapes are quality control specific. 1. Circles for quality control with trimming, 2. Triangular for sample with only trimming control and 3. Squared for sample without any quality control. Colors are distinct between the two genomes (black for *P. trichocarpa* and white for *P. tremula × P.alba*). On the left, the results for Well-Watered (WW) conditions and on the other side Water Deficit and ReWatering (WD-RW). **B**. Impact of two different tools for mapping reads, BISMARK and BSMAP on the percentage of mapped reads on two distinct genomes: *P. trichocarpa* and *P. tremula × P. alba*. Dark bars are for BISMARK and light bars for BSMAP software. On the left, the results for Well-Watered (WW) conditions and on the other side Water Deficit and ReWatering (WD-RW). Values are ‘QC’ means quality control with trimming and filtering. **C**. Percentages of cytosine considered by BISMARK after quality control and 10X minimal coverage filtering on the two genomes: *P. trichocarpa* and *P. tremula* × *P. alba*. Bars are for methylation percentage and dots are for coverage values. Black color is for Well-Watered (WW) conditions and white for Water Deficit and Re-Watering one (WD-RW). **D**. Percentages of cytosine considered by BISMARK after quality control and 10X minimal coverage filtering for the three methylation contexts (CG, CHG and CHH) for *P. tremula* × *P. alba*.

**Supplemental Figure 2:** Phenotypic and physiological characterization of the wild type and the two RNAi-*ddm1* (*ddm1*-15, *ddm1*-23) poplar lines in control (well-watered, WW) and stress (moderate water deficit followed by rewatering, WD-RW) treatments. Open symbols and open bars for WW; closed symbols and closed bars for WD-RW. Values are genotypic means ± SE. **A**. Predawn and minimum leaf water potential measured before rewatering (*n* = 5 per line per treatment). **B**. Bulk leaf carbon isotope composition (δ^13^C) measured on leaves sampled at the end of the experiment (*n* = 6 per line in WW, *n* = 12 per line in WD-RW). **C**. Total stomatal density measured at the end of the experiment (*n* = 3 per line per treatment). Treatment effects were evaluated within each line using a t-test. Levels of significance are *, 0.01 < *P* < 0.05; **, 0.001 < *P* < 0.01; ***, *P* < 0.001; ns, non-significant.

**Supplemental Figure 3: A**. Variation among *Populus* RNAi *ddm1* lines concerning the effect of leaf position on the stem on leaf area. White dots for control line and black dots for RNAi lines. Values are line means (± SE, n=3 by lines WW). For each line the rank effect was evaluated using a T-test (IF: leaf index). **B**. Total leaf area variations among Populus RNAi ddm1 lines. White bars represent well-watered conditions. Values are line mean (± SE, n=3 by lines WW). Global line effect was evaluated using a T-test (L: line). Levels of significance are *: p< 0.05, **: p< 0.01, ***: p< 0.001 and ns: non-significant

**Supplemental Figure 4:** Particular leaf phenotypes among *Populus* RNAi ddm1 lines. Open symbols for control (well-watered, WW) condition; closed symbols for stress (moderate water deficit followed by rewatering, WD-RW) condition. Values are genotypic means ± SE (*n* = 6 per line for WW and WD-RW). Treatment effects were evaluated within each line by using a t-test. The different letters indicate the differences between lines within a water regime following a Tukey’s post hoc test (small letter, well-watered condition and capital letter water deficit follow by rewatering condition). Levels of significance are *: p< 0.05, **: p< 0.01, ***: p< 0.001 and ns: non-significant.

**Supplemental Figure 5:** Variations in DNA methylation among RNAi and WT lines in shoot apical meristem one week after rewatering (t2). **A.** Global DNA methylation percentage calculated by HPLC in WT and RNAi lines in both Well-Watered (WW, white bars) and water-deficit followed by ReWatering conditions (WD-RW, black bars). Values are genotypic mean (± SE, n=3 by lines in WW and n=3 by lines in WD-RW). Global genetic variations and the effect of water deficit re-watering were evaluated using ANOVA. Different letters indicate the differences between genotypes within WW and WD-RW conditions following a Tukey’s post hoc test. Levels of significance are *: p< 0.05, **: p< 0.01, ***: p< 0.001 and ns: non-significant. **B**. Differentially methylated regions (DMRs) between treatment and lines. Black bars for hypomethylated DMRs and white bars for hypermethylated DMRs. DMRs represented are mapped against *Populus tremula* x *alba* reference genome and have passed 10X of coverage and show a cut-off of methylation of 25 % with qvalue 0.01. **C**. DMRs count between ddm1_15 and WT line. **D**. DMRs count between ddm1_23 the WT line. **E**. Distributions of the DMRs between the RNAi and the WT lines in the different contexts of methylation.

**Supplemental figure 6:** Identification of differentially methylated genes. **A**. Annotation of the common DMRs between the RNAi and the WT lines in WW and WD-RW regimes. Gene annotation was retrieved from *P. tremula* x *P. alba* v1.1 annotation. Promoters correspond to +/- 2kb from TSS (transcription start site). TE (Transposable element) annotations were retrieved from *P. trichocarpa* annotation and blasted against *P. tremula* x P. reference genome. **B**. Identification of common differentially methylated genes (DMGs) between the two RNAi lines vs. the WT line in both WW and WD-RW regimes. In DMRs = Genes that overlapped with the common DMRs, 2 kb = Genes in +/- 2 kb of DMR, 5 kb = Genes in +/- 5 kb of DMR, 10 kb = Genes in +/- 10 kb of DMR and 25 kb = Genes that are found in +/- 25 kb of DMR. **C**. Genes Ontology (GO) annotation of the common DMRs in WD-RW regimes for all contexts. GO labels were retrieved from A. thaliana annotation and a treemap was realized using REVIGO software.

**Supplemental figure 7:** Relationship between variation in DNA methylation and gene expression. **A**. Overlap between DEGs and DMRs. DEGs that are localized in DMRs or in the proximity of DMRs are represented: In DMRs = DEGs that overlapped with the DMRs, 2 kb = DEGs in +/- 2 kb of DMR, 5 kb = DEGs in +/- 5 kb of DMR, 10 kb = DEGs in +/- 10 kb of DMR and 25 kb = DEGs that are found in +/- 25 kb of DMR. **B**. Covariation between DNA methylation and gene expression. DEGs found in or near +/- 10 kb of DMRs were used for correlation (spearman t test correlation, pvalue = 0.0004).

**Supplemental figure 8:** Transposable elements activity among RNAi and WT lines in shoot apical meristem collected one week after rewatering (t2). **A**. Counts of transposable element families found in the mobilome-seq analysis in the three different lines and in WW and WD-RW regimes. White bars for WW and black bars for WD-RW. **B**. GO annotation of biological process of genes found in or near +/- 25 kb of mobilome-seq TE families. The treemap was realized using REVIGO. **C**. Overlap between TE families vs. DMRs and Genes. In = TEs that overlapped with the DMRs, 2 kb = TEs in +/- 2 kb of DMR, 5 kb = TEs in +/- 5 kb of DMR, 10 kb = TEs in +/- 10 kb of DMR and 25 kb = TEs that are found in +/- 25 kb of DMR.

**Supplemental Table 1:** Mean methylation level in ddm1 and WT lines in both WW and WD-RW conditions, Values are the average of methylation (in %) for each lines in the three different contexts (CG, CHG and CHH).

